# Mutations in *mfd* cause *Staphylococcus aureus* mucoid hyper-biofilm phenotype in chronic rhinosinusitis

**DOI:** 10.64898/2026.06.30.735446

**Authors:** Ghais Houtak, Ian R. Monk, Muhammed Awad, Roshan Nepal, Mahnaz Ramezanpour, Alkis James Psaltis, Peter-John Wormald, George Bouras, Timothy P. Stinear, Sarah Vreugde

## Abstract

Chronic Rhinosinusitis (CRS) is a common chronic inflammation of the paranasal sinus mucosa. *Staphylococcus aureus* contributes to its severity through biofilm formation. In this study, we isolated eight sequential methicillin-resistant *S. aureus* (MRSA) isolates from a patient with severe CRS over a period of 672 days (T1-T8). The isolates were phenotypically and genomically characterised, and the extracellular biofilm proteome analysed. We identified an accumulation of mutations that included the acquisition of an IS21 family insertion sequence inactivating the *icaR* gene and nucleotide variants in various genes including the transcription repair coupling factor (*mfd)*. The genomic changes were associated with a switch to a mucoid phenotype from T3 onwards (Day 178), with a significant increase in biofilm-forming capacity and the secretion of multiple enterotoxins. Targeted mutagenesis confirmed *mfd* is a regulator of strain mucoidy with enhanced biofilm and enterotoxin production. These findings support mfd as a target for novel anti-virulence therapies.

## Introduction

Chronic rhinosinusitis (CRS) is one of the most common diseases in the Western world, affecting approximately 10% of the population (Hastan et al., 2011). Although CRS is a heterogeneous disease, the primary manifestation is a chronic inflammation of the nasal and paranasal mucosal lining (Fokkens et al., 2020). *Staphylococcus aureus* is one of the most commonly isolated bacterial species from the sinuses of CRS patients (Okifo et al., 2022) and is thought to play an aggravating role in a subpopulation of CRS with nasal polyp (CRSwNP) patients (Tomassen et al., 2016; Vickery et al., 2019). Moreover, bacterial biofilm formation is associated with increased disease severity, the persistence of postoperative symptoms, ongoing mucosal inflammation and recurrent infections (Singhal et al., 2010; Zhang et al., 2011). The biofilm mode of growth protects *S. aureus* from the host immune response and increases the tolerance to antibiotics, thereby promoting persistent colonisation and recurrent infections (Flemming et al., 2016; Hall and Mah, 2017). Mucoid *S. aureus* isolates exhibit an enhanced ability to form biofilms (Jefferson et al., 2003) and have been directly isolated from sputum samples of cystic fibrosis patients (Lennartz et al., 2019; Schwartbeck et al., 2016). Mucoid *S. aureus* strains are more resistant to opsonophagocytic killing by neutrophils and survived significantly better under nutrient-limited conditions than non-mucoid isolates (Schwartbeck *et al*., 2016). A specific 5-bp deletion in the *icaR-icaA* intergenic region can cause a high biofilm-forming mucoid phenotype (Brooks and Jefferson, 2014). The *icaADBC* operon encodes enzymes involved in the synthesis of polysaccharide intercellular adhesin (PIA), a cell surface-associated exopolysaccharide, also called poly-β(1-6)-N-acetylglucosamine (PNAG), and the primary extracellular polymeric substance (EPS) of *S. aureus* biofilms (Arciola et al., 2015; Rohde et al., 2007). PIA/PNAG mediates bacterial adhesion, leading to intercellular aggregation and biofilm accumulation (Haaber et al., 2012; Mack et al., 1996).

Although within-host single nucleotide polymorphisms (SNP) occur at a low rate (∼8 mutations per genome per year) in *S. aureus* (Didelot et al., 2016), adaptive protein-altering mutations have been demonstrated to occur in metabolic genes, in regulators of quorum-sensing and in known antibiotic targets (Coll et al., 2025; Giulieri et al., 2022). Further, SNPs in regulatory genes have been associated with significant phenotypic changes, including biofilm formation, virulence and pathogenicity (Sharkey Liam et al., 2023; Viana et al., 2015). Additionally, the gain or loss of mobile genetic elements (MGEs) like prophages, plasmids, and genomic islands have been linked with human-host adaptations (Chaguza et al., 2022; Houtak et al., 2023). *In vitro* experiments have shown that genetic alterations promoting adaptive evolution in bacteria can develop in less than 500 generations (∼30 days) (Bennett and Hughes, 2009).

In this study, we assembled closed genomes of eight longitudinally isolated (over 672 days) clones of *S. aureus* and characterised phenotypic parameters (growth kinetics, biofilm production, extracellular proteome) to identify the consequence of the mucoid phenotype. Our results reveal within-host adaptation of a persistent *S. aureus* isolate towards a mucoid hyper biofilm-forming isolate with increased auto-aggregation and virulence factor secretion associated with chronic sinus inflammation. This study offers a new perspective on the bacterial factors contributing to hyper biofilm-forming mucoid *S. aureus* phenotypes in chronic inflammatory disease.

## Materials and methods

### Ethics

This study complies with all relevant ethical regulations for human research. Ethical approval was obtained from the Central Adelaide Local Health Network Human Research Ethics Committee under the following reference number: Ref no. 13604.

### Bacterial isolates

Clinical Isolates (CIs) were retrieved from our bacterial biobank comprised of samples collected from ear-nose-throat inpatient and outpatient clinics. All the *S. aureus* clinical isolates used in this study (N=8) were isolated from a single CRS patient at multiple time points. For all experiments, the CIs were cultured at 37°C overnight from glycerol stocks onto nutrient agar (NA) plates (Thermo Fisher Scientific, CM0003, Waltham, USA) unless specified differently. To distinguish mucoid phenotypes of *S. aureus*, clinical isolates were cultured on Congo Red Agar (CRA). The CRA was prepared using Brain Heart Infusion Agar (Thermo Fisher Scientific) with the addition of 0.08% Congo Red (Sigma-Aldrich, C6767-25G, St Louis, USA) and 5% sucrose (w/v) as specified by Freeman et al. (Freeman et al., 1989). The criteria for determining mucoid colonies among the CIs involved examining their colony morphology on CRA. Mucoid colonies were identified as dry with irregular edges, while non-mucoid colonies were characterized by their smooth and circular appearance as specified by Schwartbeck et al. (Schwartbeck *et al*., 2016).

### *S. aureus* growth curve analysis

The growth parameters of all CIs were assessed by measuring the absorbance of broth cultures at 600 nm (OD_600_) using a SmartSpec 3000 UV/Vis spectrophotometer (Bio-Rad Laboratories Inc., Hercules, USA). Growth assays were performed in triplicate. Briefly, a few colonies from overnight nutrient agar (NA) plates were resuspended in sterile saline to obtain 1.0 McFarland standard unit (MFU). One hundred microliters of the bacterial suspension was added to 15 ml of Tryptic Soy Broth (TSB) in a 50 ml Falcon tube. The tubes were incubated at 37°C in a shaking incubator (180 rpm). Every hour, 100 µL of culture was pipetted and mixed with 900 µL sterile broth in a cuvette for OD reading till 8-hours post-inoculation. The last reading was taken after 24 hours post-inoculation. The growth curves were analysed using the R package Growthcurver v0.3.1 (Sprouffske and Wagner, 2016).

### Determination of biofilm biomass

Biofilm biomass was determined using the Crystal Violet (CV) assay with biofilms established for 48 hours in 96-well flat-bottomed tissue culture-treated microtiter plates (Corning, 3599, Corning, USA). Briefly, the overnight broth cultures of CIs were adjusted to 1.0 McFarland and diluted 1:15 in growth media. One hundred fifty microliters of each diluted culture were pipetted in inner wells of 96-well plates and incubated at 37°C on an orbital shaker (80 rpm). After 48 hours of incubation, planktonic cells were carefully aspirated, and the plates were washed twice with Phosphate Buffer Saline (PBS, 1X). Plates were air-dried, and 200 ml of 0.01% CV solution was added to each well and left at room temperature for staining. After 10 minutes, the excess CV was aspirated, washed thrice with saline, and air-dried. Finally, the biomass-bound crystal violet was solubilised in 200 ml of 30% acetic acid. The biomass was monitored in terms of absorbance (OD_600_) at 600 nm using CLARIOstar Plus (BMG Labtech, Ortenberg, Germany). Three different media, nutrient broth (Thermo Fisher Scientific, CM0003), Mueller-Hinton broth (Thermo Fisher Scientific, CM0405), and tryptic soy broth (Thermo Fisher Scientific, CM0129), were used to assess the biofilm in different nutrient conditions. The crystal violet assay was performed in triplicate with six technical replicates. RN4220 (ATCC, Manassas, USA), a potent PIA-dependent biofilm-forming laboratory strain, was used as a reference strain (Nair et al., 2011; O’Neill et al., 2008).

### Determination of antibiotic sensitivity

The sensitivity of *S. aureus* isolates to mupirocin, levofloxacin, amoxicillin/clavulanic acid and vancomycin (Sigma-Aldrich, St Louis, USA) was determined using broth microdilution assay (Wiegand et al., 2008). Single colonies of freshly streaked plates were collected, and bacterial suspensions were adjusted to 0.5 MFU, followed by 1:100 dilution in Mueller Hinton broth. Two-fold dilutions of each antibiotic were prepared in the same media. The minimum inhibitory concentration (MIC) of each antibiotic was determined as the minimum concentration that inhibits bacterial growth.

### Bacterial genomic DNA extraction and sequencing

Genomic DNA (gDNA) was extracted using 1 mL (from 10 mL overnight TSB culture) *S. aureus* culture using the DNeasy Blood & Tissue Kit (Qiagen, 69504, Hilden, Germany) according to the manufacturer’s guidelines (Supplementary text ST1). The extracted gDNA was sequenced using long-read Oxford Nanopore Technology (ONT) on the MinION Mk1C (Oxford Nanopore Technologies, Oxford, UK) using R9.4.1 MinION flowcells (Oxford Nanopore Technology) with the Rapid Barcoding Kit (Oxford Nanopore Technology, SQK-RBK 110.96), according to the manufacturer’s instructions using 50 ng of isolated gDNA. Base-calling was conducted with Guppy v 6.2.11 super accuracy mode using the ‘dna_r9.4.1_450bps_sup.cfg’ configuration (Oxford Nanopore Technology). *S. aureus* isolates were also sequenced using the NextSeq 500/550 Mid-Output sequencing kit v2.5 and the NextSeq 550 Illumina platform (Illumina Inc, San Diego, USA) and NextSeq 500/550 Mid-Output sequencing kit v2.5. The sequencing was conducted at a commercial facility (SA Pathology, Adelaide, SA, Australia) as previously described (Shaghayegh et al., 2023). In short, gDNA was isolated using the NucleoSpin Microbial DNA kit (Machery-Nagel GmbH and Co.KG, 740235.50 Duren, Germany). Sequencing libraries were prepared using a modified protocol for the Nextera XT DNA library preparation kit (Illumina Inc., FC-131-1024). The genomic DNA was fragmented, after which the amplification of Nextera XT indices to the DNA fragments was performed utilising a low-cycle PCR reaction. One hundred fifty bp reads were obtained by sequencing after purification and normalisation of the amplicon library.

### Bioinformatic Analysis

Chromosome assemblies: Complete chromosomal *S. aureus* assemblies were created using Hybracter v0.4.0 (Bouras et al., 2024b). Briefly, long-read adapters and barcodes were trimmed using Porechop v0.2.4 (Wick, Porechop). Short reads were filtered, with low-quality regions and adapters trimmed using fastp v0.23 (Chen et al., 2018). Long-read-only assemblies were created using Flye v2.9.2 with ‘—nano-hq’ specified (Kolmogorov et al., 2019). Assemblies, including contigs with a length greater than 2.5 Mb, were kept and denoted as the putative chromosomes. Targeted plasmid assembly was then conducted using Plassembler v1.4.1 (Bouras et al., 2023b). The resulting chromosomes were polished with long reads first using Medaka v 1.8.0 (Oxford Nanopore Technologies, 2022) using the r941_min_sup_g507 model. After the first round of polishing, the chromosomes were reoriented to begin at the *dnaA* gene using Dnaapler v0.4.0 (Bouras et al., 2024a), then polished again with Medaka and subsequently with short reads using Polypolish v 0.5.0. (Wick and Holt, 2022), followed by pypolca v0.2.0 (Bouras et al., 2024c) a standalone Python reimplementation of POLCA (Zimin and Salzberg, 2020).

### Genomic analysis

Finished *S. aureus* genome assemblies were annotated with Bakta v 1.5.0 (Schwengers et al., 2021). All CIs were genotyped (clonal complex) according to the PubMLST scheme (Jolley et al., 2018; Seemann, mlst). Antimicrobial resistance and virulence genes were identified by screening genome sequences through the Comprehensive Antibiotic Resistance Database (Jia et al., 2017) and Virulence Factor Database (Liu et al., 2019) using ABRicate, v 1.0.1 (Seemann, Abricate). To detect structural changes such as DNA sequence repeats, large deletions, or insertions, we used NucDiff v2.0. (Khelik et al., 2017). We used Snippy to detect single nucleotide polymorphisms (SNPs) between CIs (Seemann, 2015). Dot plots were generated using D-GENIES v1.4 performing large genome alignments using minimap2 (Cabanettes and Klopp, 2018). Prophage sequences were detected by running hlbroken on the CI genomes (Bouras, hlbroken). Resultant prophage genomes were annotated using PhiSpy v4.2.21 (Akhter et al., 2012), Pharokka v 1.5.0 (Bouras et al., 2023a) and Phold v1.0.0 (Bouras et al., 2026).

Protein structure prediction was conducted using the Colabfold webserver v1.5.3 (Mirdita et al., 2022). Models were generated without relaxation and the top ranked model in terms of pLDDT was retained. Predicted structures were queried against the Alphafold Database (Letunic et al., 2021) using the Foldseek webserver (using v8-ef4e960) (van Kempen et al., 2024). Predicted Aligned Error and structures were visualised with the PAE Viewer webserver (Elfmann and Stulke, 2023).

### Secretome extraction, digestion, and sample preparation for DIA-MS

Data-independent acquisition mass spectrometry of growth culture supernatants was performed for T1, T3, T5, T6, and T8. Single colonies from overnight NA plates were dissolved in sterile saline to obtain 1.0 MFU. One hundred microliters of the solution were added to 15.0 ml of TSB and incubated at 37 °C in a shaking incubator (180 rpm). After 7 hours, the tube was briefly vortexed and centrifuged at 4000 x g for 10 min. The secretome was sterilised by passing through a 0.22 µm acrodisc filter (Pall Corporation, 4612, New York, USA) and concentrated using a 3K MWCO Pierce Protein Concentrator PES (Thermo Fisher Scientific, 88525). The protein concentration was determined using NanoOrange Protein Quantitation Kit (Thermo Fisher Scientific, N6666) as per the manufacturer’s instruction. The proteomics of the secretome were analysed using a data-independent acquisition mass spectrometry (DIA-MS) using Orbitrap Fusion Lumos Tribrid Mass Spectrometer (Thermo Fisher Scientific) with Dionex Ultimate 3000 UPLC (ThermoFisher Scientific) according to an established protocol. Detailed settings of the runs can be found in Supplementary text ST2.

### Protein identification, quantification, and DIA data analysis

The DIA spectra were processed and quantified using Spectronaut v15 (Biognosis AG, Schlieren, Switzerland) with factory default settings. An in-house *S. aureus* proteome database created from all genes identified in one of the longitudinal clinical isolates (T6) was used as a reference. Biological processes and gene annotations were assigned based on the previously annotated *S. aureus* T6 isolate using Bakta (Schwengers *et al*., 2021).

Differential protein expression analysis was performed in R v4.2.0 (R Core Team, 2017) using the DEP package v1.20.0 to calculate differentially expressed proteins (DEP) (Zhang et al., 2018). Functions from the tidyverse collection of R packages v 1.3.2 were incorporated into the analysis and visualisation (Wickham et al. 2019). The threshold for identifying differentially expressed proteins was set at a false discovery rate (FDR) of less than 0.05.

### Construction of *S. aureus* mutants

Isolate T1 encodes a full-length Mfd protein (1168 amino acid), compared to a truncated 763 amino acid variant encoded by T3 starting from the N-terminal end, along with a 419 amino acid frameshifted open reading frame constituting the C-terminal end of the Mfd protein. This arises from the deletion of a directly repeated “ATTAAAAC” sequence in *mfd* gene of the T3 isolate. As a barrier to transformation, isolate T1 contains three type I *hsdS* alleles with one of them equivalent to the SCC*mec* adjacent allele (SAA6159_00055) of *S. aureus* strain JKD6159. SAA6159_00055 and its cognate methylase (SAA6159_00054) were previously introduced into the chromosome of *E. coli* DC10B (DH10BΔ*dcm*) yielding SA93Bv1.0 (Monk et al., 2015). T1 also contains a putative type IIG system with the recognition sequence of “YAC(N5)TGG” (Roberts et al., 2022). To facilitate bypass of the type IIG system, the putative type IIG methylase was amplified with primers (IM1779/IM1780), digested (NcoI/SalI) and ligated into the *E. coli* replicating plasmid pIMK4 under the control of a weakly expressed constitutive promoter (Monk et al., 2008). The ligation was then transformed into SA93Bv1.0 yielding SA93Bv1.0+pIMK4 (TIIG).

Primers were designed to amplify ∼500 bp upstream (IM1769) and downstream (IM1770) of the deletion from T3 and cloned into the pIMAY-Z allelic exchange vector as described previously (Monk and Stinear, 2021) and transformed into SA93Bv1.0+pIMK4(TIIG) yielding plasmid pIMAY-Z(*mfd^-^T3*). Purified pIMAY-Z(*mfd*^-^*T3*) was transformed into isolate T1 and the transformants were selected on brain heart infusion agar containing 10 µg/ml of chloramphenicol and 100 µg/ml of X-gal. Only two transformants were obtained from 3 independent electroporations. The allelic exchange protocol of Monk and Stinear (Monk and Stinear, 2021) was followed with the white colonies from plasmid excision screened on Congo Red agar for a smooth to rough transition with comparison to T1 (smooth) and T3 (rough). This yielded isolate T1*^mfd-^*^T3^. Genomic DNA was isolated from T1*^mfd-^*^T3^ and sequenced on the ONT platform (flowcell R10.4.1) through the Centre for Pathogen Genomics, University of Melbourne.

IM1769mfd/F:CCTCACTAAAGGGAACAAAAGCTGGGTACCATAAAGAAAGAGAAATGGCAGAAGG IM1770mfd/R:CGACTCACTATAGGGCGAATTGGAGCTCTGTAGGAAGTATGCATAACCAATACG IM1779 C222 (NcoI) typeIIGm/F atatCCATGGacgaaaaaactggtatagacc IM1780 C222 (SalI) typeIIGm/R atatGTCGACttataatctatctaatatcttattttcttttcgcttc Further, multiple mutants in the standard reference strain JE2 (CA-MRSA USA300 clone, ATCC, Manassas, USA) were constructed to elucidate the roles of *icaR* and *mfd* genes alone or in combination for the development of the hyper-biofilm forming mucoid phenotype. The mutants were then cultured on CRA for phenotypic assessment (mucoid vs non-mucoid) and biofilm biomass was estimated as described before.

Minimap2 v2.28 (Li, 2018), Samtools v1.21 (Li et al., 2009) and IGV v 2.15 (Robinson et al., 2011) were used to map, visually inspect and confirm the mutants possessed the desired variants.

### Statistical analysis

Statistical analysis was conducted using R v4.2.0 (R Core Team, 2017). The data was presented as the mean ± standard error of the mean (s.e.m.). Significance was determined at a p-value of less than 0.05 after correction for multiple comparisons.

### Data availability

The *S. aureus* whole genome assemblies and short and long read FASTQ files generated in this study are publicly accessible on the Sequence Read Archive (SRA) under the BioProject accession PRJNA914892. The following BioSample accessions correspond to the samples analysed: T1 (SAMN32360844), T2 (SAMN32360968), T3 (SAMN32360969), T4 (SAMN32360970), T5 (SAMN32538168), T6 (SAMN32360890), T7 (SAMN32360971), and T8 (SAMN32360972). The *S. aureus* mutant long read FASTQ files generated in this study are publicly accessible on the Sequence Read Archive (SRA) under the BioProject accession PRJNA1050835. The following BioSample accessions correspond to the samples analysed: C1137 (SAMN38755502), C1138 (SAMN38755503). The protein structures (.pdb) with associated scores (.json) predicted using Colabfold for the T1 full-length Mfd protein and the truncated T3-T8 Mfd protein can be found as Supplementary data.

### Code availability

All code used in this manuscript can be found at https://github.com/gbouras13/Houtak_et_al_longitudinal_Staphylococcus_aureus.

## Results

### *S. aureus* isolates from a severe recalcitrant CRS patient evolved into a mucoid phenotype with the propensity to self-aggregate

Methicillin-resistant *S. aureus* (MRSA) clinical isolates (n=8) were longitudinally isolated from the sinonasal cavities of a 70-year-old male diagnosed with chronic rhinosinusitis with nasal polyps (CRSwNP). The first isolate (T1) was obtained 08/01/2014 and the last (T8) 11/11/2015. The total time between the collection of the first (T1) and the last (T8) isolate was 672 days. At the time of T1 collection, the patient was had undergone eight previous sinonasal surgeries. Between the collection of T1 and T8, the patient had received multiple oral antibiotic courses, including doxycycline, amoxicillin/clavulanic acid, ciprofloxacin, and trimethoprim/sulfamethoxazole (Figure 1). Courses of topical antibiotics were also given occasionally in the form of nasal rinses with mupirocin or tobramycin. Furthermore, the patient had received topical corticosteroids (budesonide nasal rinses) during the majority of the time between the collection of T1 and T8 and two courses of oral corticosteroids. Complete genome sequences were established for all eight isolates including 2 isolates (C222 and C333) that had been sequenced previously (Houtak et al., 2023). A summary of the isolation dates and general genomic characteristics of the isolates is provided (Table 1).

**Figure 1.**
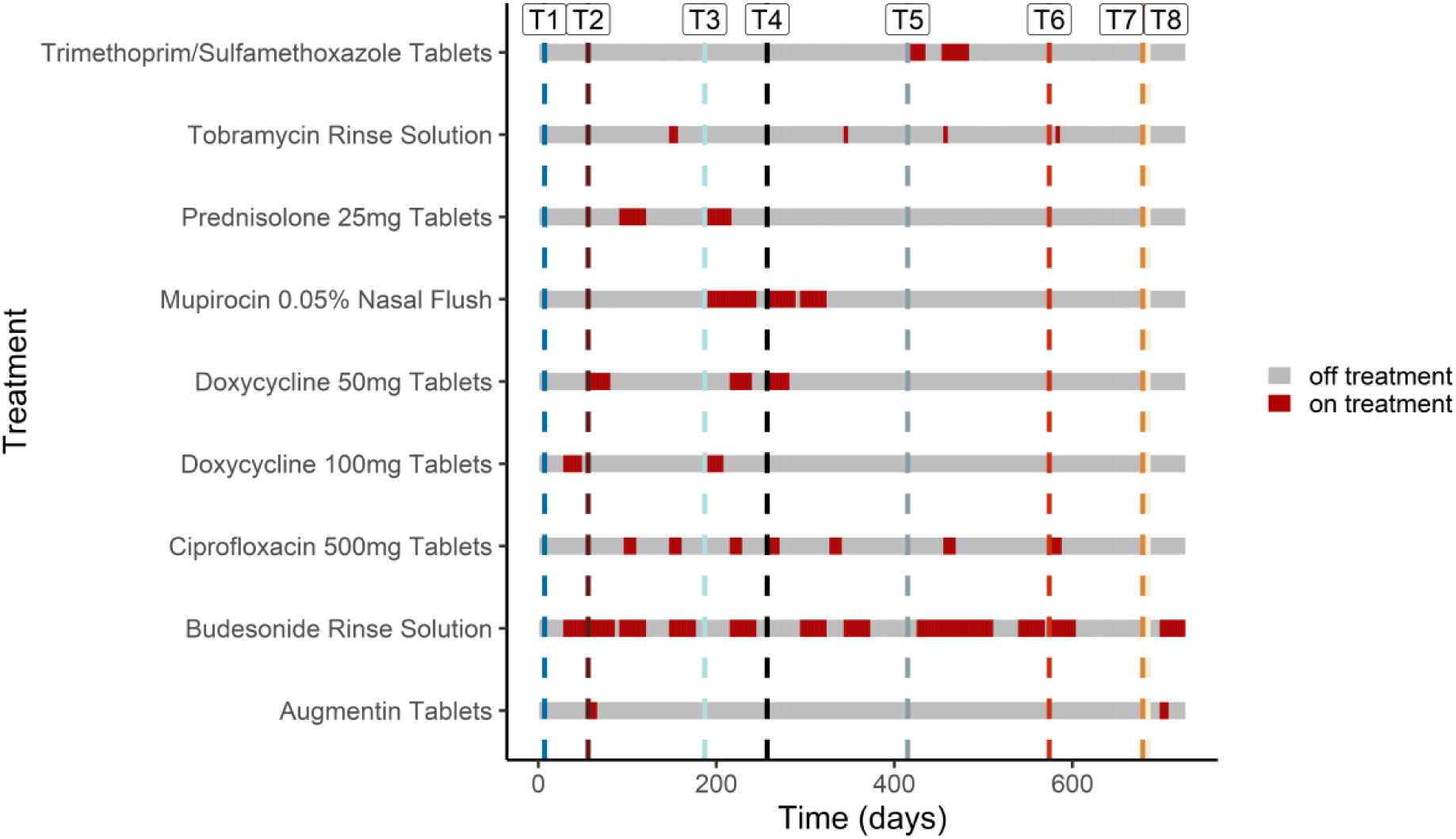
Timeline of the patients’ therapeutic regimen with collection points of the 8 *S. aureus* isolates. Vertical dashed lines indicate the time of collection.

**Table 1.** Details of clinical isolates of Staphylococcus aureus used in the study.

| **Bacterial isolate\*** | **Collection date** | **Genome size (bp)** | **Days between collection** | **Clonal complex / Sequence type** |
| --- | --- | --- | --- | --- |
| **T1 (C222)** | 08/01/2014 | 2,788,579 | 0 | CC22/ST22 |
| **T2 (C229)** | 26/02/2014 | 2,857,176 | 49 | CC30/ST30 |
| **T3 (C247)** | 07/07/2014 | 2,834,886 | 180 | CC22/ST22 |
| **T4 (C272)** | 15/09/2014 | 2,834,854 | 250 | CC22/ST22 |
| **T5 (C308)** | 20/02/2015 | 2,834,886 | 408 | CC22/ST22 |
| **T6 (C333)** | 29/07/2015 | 2,837,533 | 567 | CC22/ST22 |
| **T7 (C344)** | 21/10/2015 | 2,835,329 | 651 | CC22/ST22 |
| **T8 (C347)** | 11/11/2015 | 2,837,534 | 672 | CC22/ST22 |

\* The first isolate is termed T1, and later isolates are termed T2-T8 (reference clinical isolate code in NCBI genome database are between the parentheses). CC= clonal complex, ST= sequence type.

We screened the eight isolates for differences in biofilm biomass when grown in three different growth media. All isolates except T2 had higher biofilm biomass than a strong biofilm-forming reference strain RN4220 across the different media (p<0.05), indicating the ability of these isolates to form robust biofilms (Sugimoto et al., 2018). Isolate T1 consistently yielded a lower biofilm biomass compared to isolates T3-T8 in MHB and TSB media (p<0.05) (Figure 2). When normalised to the biomass of RN4220, the largest biomass ratios between the clinical isolates and RN4220 were seen in MHB, where the ratio between T1 and RN4220 was 1.95. In contrast, the ratio between T2 and RN4220 in MHB was 0.3. However, the ratios between the later timepoint isolates to RN4220 were consistently higher, between 3.96 and 4.93 (Table S1).

**Figure 2.**
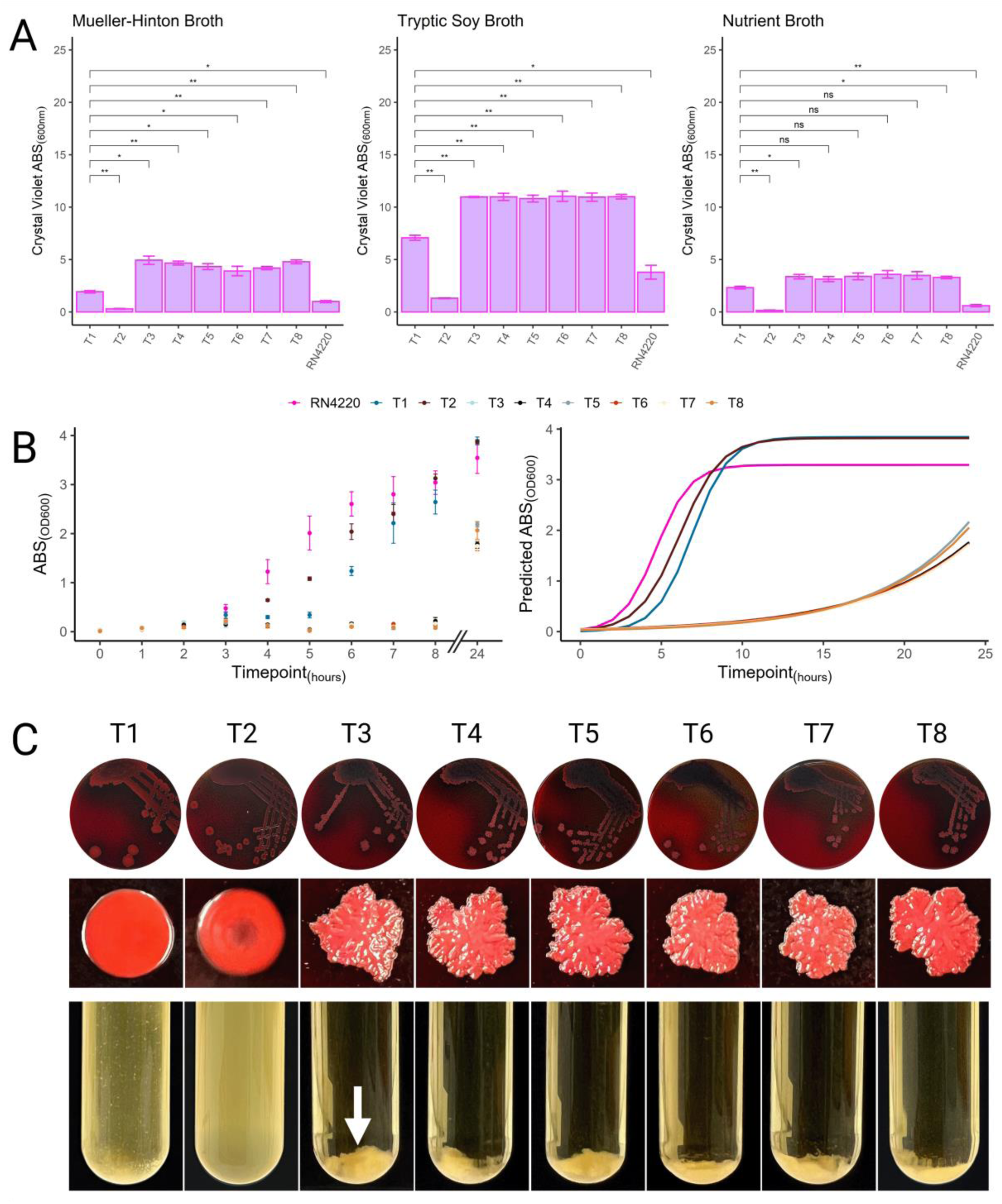
Analysis of *S. aureus* isolates collected from a CRS patient with severe disease. (A) Biofilm quantification of the 8 consecutive clinical isolates and RN4220 using the crystal violet assay in various media. (B) OD600nm measurements in tryptic soy broth for all isolates and RN4220, with predicted growth curves shown on the right. (C) Congo red agar plates, zoomed-in images of single colonies, and broth culture turbidity after 6 hours for all isolates, with aggregated bacterial cell sedimentation indicated by an arrow. Statistical significance was determined using one-way ANOVA and pairwise post-hoc T-test with Benjamini-Hochberg p-value adjustment (*p<0.05, **p<0.01).

When the isolates were cultured on CRA, all produced dark colonies, signifying the production of PIA/PNAG. All isolates except for T1 and T2, showed a mucoid phenotype, as depicted in Figure 2. When cultured on TSA, the mucoid isolates T3-T8 presented a highly viscous and sticky phenotype, resulting in pronounced adhesion between cells (Figure S1). The most striking phenotypic difference between the isolates was observed when cultured in TSB. During growth in TSB, the mucoid isolates (T3-T8) exhibited a dramatic increase in bacterial cell auto-aggregation during exponential growth resulting in a loss of turbidity and the settling of clumped cells at the bottom of the culture tube (Figure 2C). The T2 isolate showed no observable formation of macroscopic clumps. While the T1 culture exhibited some macroscopic auto-aggregation, it was less pronounced than the mucoid isolates T3-T8 (Figure 2C). The turbidity of the mucoid isolates increased in the stationary phase of growth and was characterised by a combination of aggregated and non-aggregated cells but did not reach the same level as the non-mucoid isolates. These results suggest that the increase in biofilm biomass between T1 and any of T3-T8 mucoid *S. aureus* isolates is potentially due to increased auto-aggregation of bacterial cells.

### Genomic comparison of the longitudinal isolates indicated mutations in the *icaR*-*icaADBC* operon were associated with a hyper-biofilm-forming mucoid phenotype

We next compared the complete genome sequences of the eight isolates. Genome length varied from 2,788,579 bp for T1 to 2,837,534 bp for T8 (Table 1). T1 and T3-T8 belonged to clonal complex 22 (CC22) while T2 was CC30. Pairwise genome alignments of all CIs to the T1 chromosome revealed that for T2, more than 50% of the genomic alignment had less than 75% alignment identity. In contrast, T1 to T3-T8 showed a near-identical pairwise alignment (>95 % alignment identity) except for the acquisition of Sa3int prophage (phiSA3) in T3-T8 near the 2 Mbp, not present in T1 (Figure 3A and Figure S2) (Nepal et al., 2021). These results suggest that T1 and T3-T8 are clones of an atypical mucoid *S. aureus* isolate that has evolved *in vivo* over the course of the patient’s treatment for CRSwNP. In contrast, the T2 isolate was a distinct genotype, suggesting the patient could have been colonised with multiple *S. aureus* lineages at that time point.

**Figure 3.**
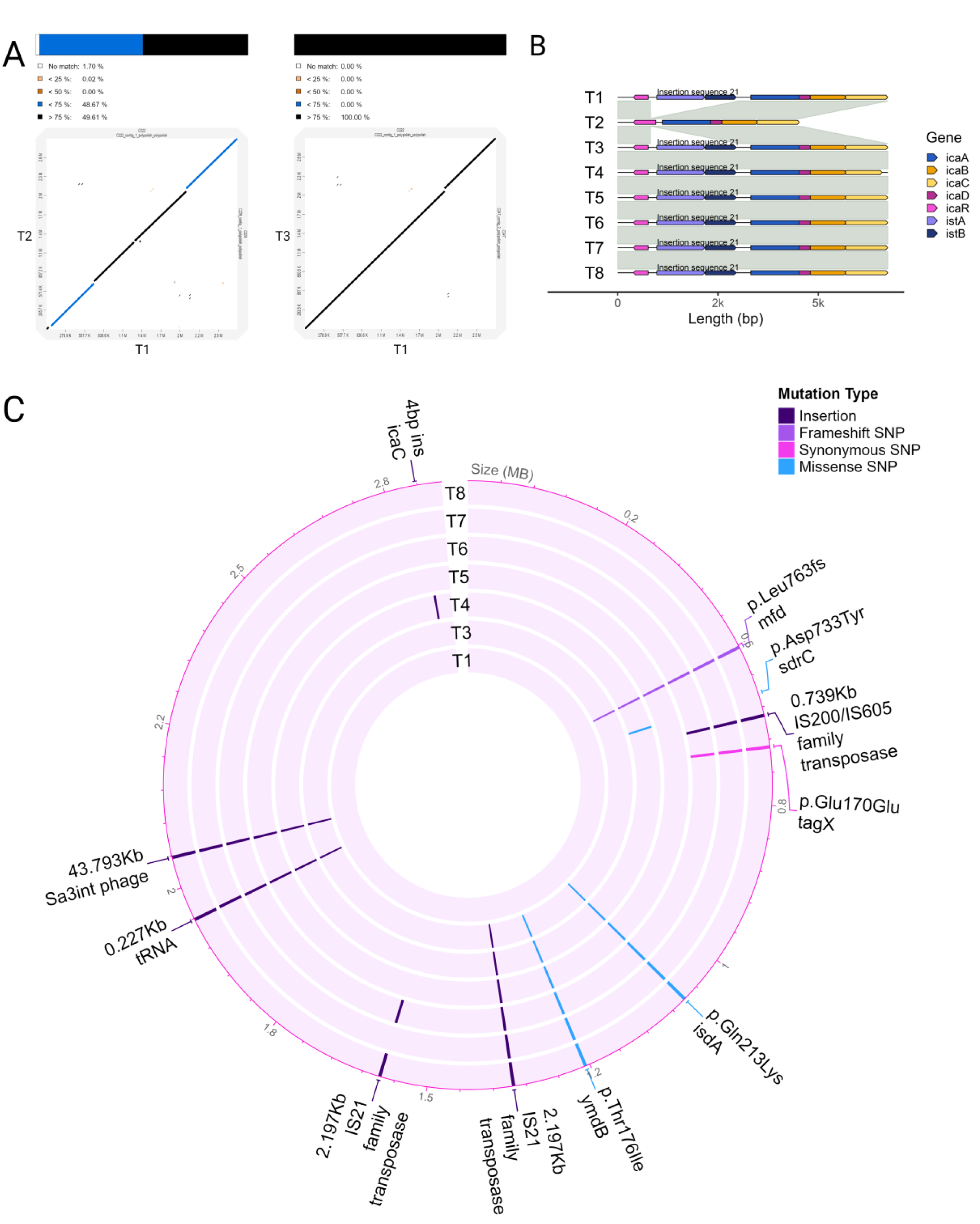
Pairwise genome alignment of the CC22 clinical isolates. (A) Dot plot comparison of T1 vs T2 and T1 vs T3, with blast-like alignment identity percentage indicated by colours. (B) Alignment of the *icaR-icaADBC* operon starting at the *icaR* gene. Synteny and sequence similarity are indicated by grey fill connecting the genomes. Genes have been highlighted in different colours. Genes belonging to the insertion sequence (IS21) are annotated. (C) The light purple rings represent the circular genome map of the closely related high biofilm-forming isolates (T1, T3-T8). Highlighted regions represent non-synonymous sequences, including prophages, insertion sequences, deletions, and single nucleotide polymorphisms, compared to T1.

To investigate the potential relationship between the *icaR-icaADBC* operon and hyper-biofilm-forming mucoid *S. aureus* isolates through increased PIA production, we compared the DNA sequence at the locus between all eight isolates. All high biofilm-forming isolates (T1, T3-T8) contained an insertion sequence (IS) belonging the IS21 family inserted into *icaR* gene, resulting in its truncation. Interestingly, read-mapping highlighted variation in the *icaC* gene region, due to the presence of a repeating 4 bp “TTTA” sequence (Figure S3). This repeat motif was mostly seen in three copies, but occasionally in four copies. The four copy variant resulted in a frameshift at residue 290, ultimately truncating the protein to length 304AA from 350AA (Figure S3C-D) and affecting the formation of biofilm in this subpopulation. The ratio of three to four copies varied between isolates. At T1, TTTA was only present in three copies, but at some timepoints (T3, T4, T7 and T8), TTTA was present in four copies in some reads (Figure S3E) and in T4, most reads contained the four-repeat motif (Figure S3).

### The genomic divergence between clinical isolates over time was determined mainly by mobile genetic elements

To identify the genetic mechanism behind the acquisition of the mucoid phenotype in T3 and beyond, we examined the genomic similarities and differences between the isolates. Our analysis revealed several structural and mutational divergences in T3 compared to T1. Whilst a Sa2int prophage was present in T1 and T3-T8, a Sa3int prophage (phiSA3), 43,793 bp long, was inserted *de novo* near the 2 Mbp location (2,044,511 bp -2,088,303 bp) in T3, disrupting the *hlb* gene. Secondly, an IS21 insertion sequence was found starting at the 1,365,841 bp in T3, leading to the disruption of an AAA family ATPase gene. Furthermore, a transfer RNA (tRNA) sequence coding for tRNA^Asp^, tRNA^Fmet^, and tRNA^Ser^ was acquired at the 1,945,485 bp in the genome of T3. Further mutational changes included SNPs and deletions in 3 genes in T3-T8. These included a deletion of 8 bp (ATTAAAAC) located in the transcription repair coupling factor Mutation Frequency Decline (*mfd*) gene, resulting in a frameshift mutation. Single SNPs resulting in missense mutations were also found in iron-regulated surface determinant protein A (*isdA*) and the metallophosphoesterase *ymdB* (Figure 3C). Analysis between the later timepoint isolates (T4-T8) and T1 showed a few additional structural and mutational divergences from the ones described for T3, such as the expansion of the 2197 bp IS21 insertion sequence in two locations (all T3-T8 gained an IS21 element around location 1.36Mbp, while T6 and T8 gained a copy at 1.57Mbp) and a SNP in the *tagX* gene.

All high biofilm-forming isolates (T1, T3-T8) contained an identical 2473 bp plasmid harbouring the *ermC* gene. Similarly, no difference in the count of antibiotic-resistance genes was observed and antibiotic susceptibility to various antibiotics tested was similar across the isolates (Supplementary Table S4). Due to the acquisition of phiSA3 in T3-T8, *scn* (staphylococcal complement inhibitor), *chp* (chemotaxis inhibitory protein) and *sak* (staphylokinase) were identified. Interestingly, T2 (the CC30 isolate) carried the most virulence factors out of all the isolates (Figure S4).

### Secretome results indicated an increased secretion of virulence factors associated with hyper-aggregation

Proteomic assessment of the potential changes in the *S. aureus* secretome across time was performed to comprehensively study the effect of the structural and mutational divergences on secreted proteins. Five isolates (T1, T3, T5, T6 and T8) were selected for analysis. A total of 1371 proteins were detected across all isolates. All comparisons were made relative to the secretome of T1. PCA separated the secretome of T1 from the other isolates (1st component 35.1% of the variance, 2nd component 18.6% of the variance). The secretome profiles of the T3, T5, T6 and T8 samples clustered together, suggesting a global similarity (Figure S5). Differential expression analysis grouped by isolate revealed differentially expressed proteins (DEPs) between all later timepoint isolates (T3, T5, T6 and T8) and T1. There were 71 DEPs between the secretome of T3 and T1 (adjusted p-value<0.05) (Figure 4A), 42 DEPs between T5 and T1 (adjusted p-value<0.05) (Figure 4B), 55 DEPs between T6 and T1 (adjusted p-value<0.05) (Figure 4C) and 43 DEPs between the secretome of T8 and T1 (adjusted p-value<0.05) (Figure 4D). Since the later timepoint isolates shared a similar phenotype and clustered together based on the PCA, we looked at consistently up- and down-regulated proteins between the secretomes of later timepoint isolates (T3, T5, T6, T8) and T1. A total of 14 proteins were significantly down-regulated, and 26 were significantly up-regulated in the secretomes of at least three later timepoint isolates compared to T1 (Figure 4E). Information on significant DEPs is shown in Table 3. Among the 14 consistently down-regulated proteins, five were encoded by the Sa2int prophage. These included the BppU-N domain-containing protein, phage portal protein, phage major capsid protein, a hypothetical phage protein, and the phage terminase small subunit P27 family protein. The Hlb protein (beta-toxin), was also consistently down-regulated, as expected with the insertion of a Sa3int prophage in the *hlb* gene resulting in a truncated protein (Nepal et al., 2025; Nepal *et al*., 2021). Furthermore, several cell-wall anchored (CWA) proteins involved in cell adhesion, such as SasG, SdrD and FnbA were also downregulated (Table 2).

**Figure 4.**
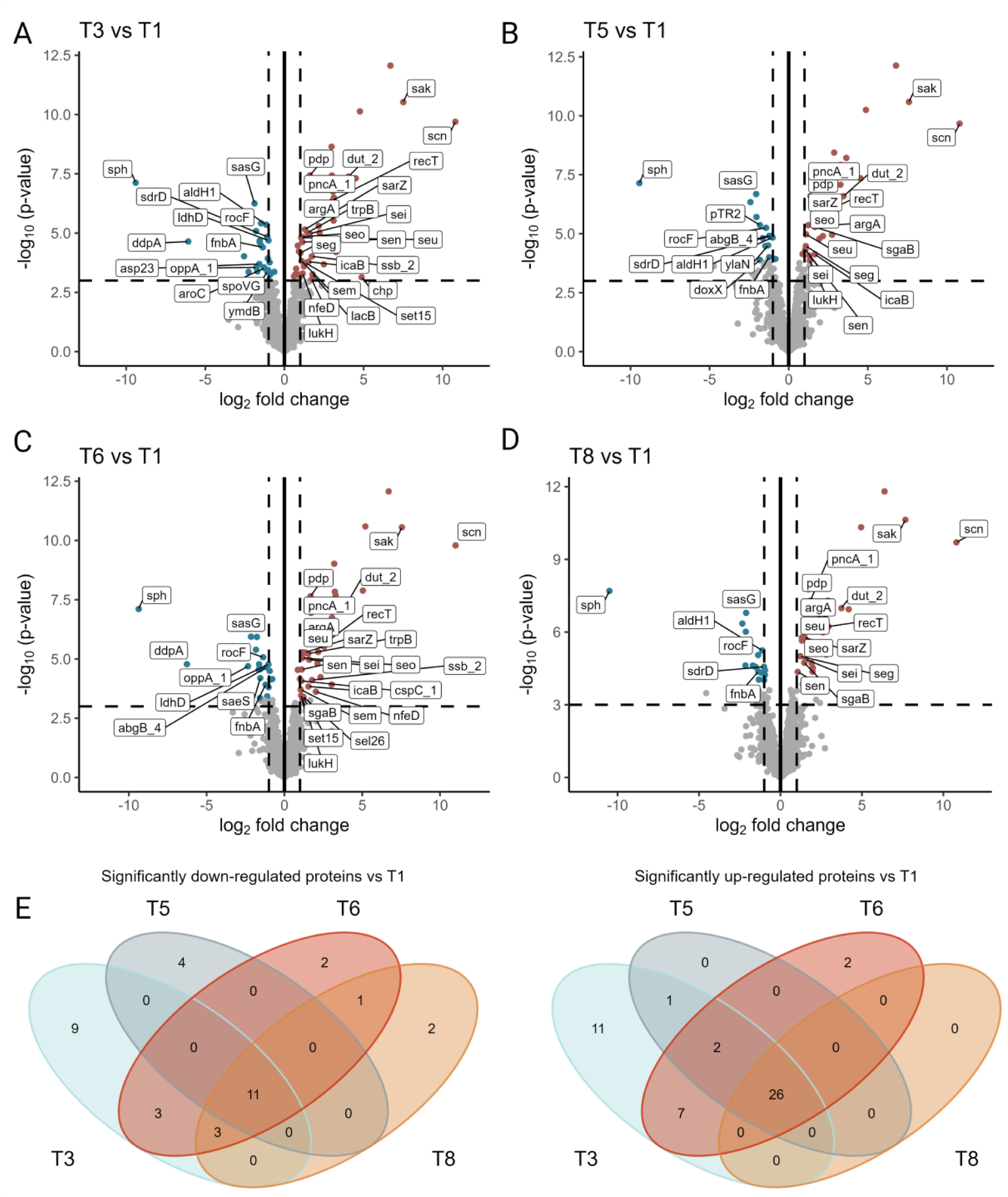
Analysis of differentially expressed proteins in the secretome of *S. aureus*. A-D volcano plots demonstrate the quantification of proteins in later timepoint isolates (T3, T5, T6, T8) compared to T1. Proteins that are significantly up-regulated (adjusted p<0.05) are represented in red and those that are significantly downregulated (adjusted p<0.05) are shown in blue. Proteins that are not differentially expressed are depicted in grey. Log2 fold change is used to express the data, with the vertical dashed lines set at -1.5 and +1.5 and the horizontal dashed line at -3 log10. (E) Venn diagram illustrates the overlap of up-and down-regulated DEPs (adjusted p-value<0.05) between the four groups (T3, T5, T6, T8).

**Table 2.** Consistent downregulation of proteins in the secretome of later timepoint isolates (T3, T5, T6, T8) compared to T1.

| Name | Product | UniRef ‡ |
| --- | --- | --- |
| **Abgb** | amidohydrolase | UniRef100_Q79EU8 |
| **AldH1** | 4,4'-diaponeurosporen-aldehyde dehydrogenases | UniRef100_UPI00128F7EEA |
| **FnbA** | fibronectin-binding protein A | UniRef90_UPI001363E29B |
| **group_104 †** | BppU-N domain-containing protein | UniRef100_I1W5Y0 |
| **group_171 †** | phage portal protein | UniRef90_M1T2X0 |
| **group_204 †** | phage major capsid protein | UniRef100_UPI0002598D2F |
| **group_294** | DUF4352 domain-containing protein | UniRef100_UPI00020F1DEF |
| **group_843 †** | hypothetical protein | UniRef100_U5U447 |
| **group_876 †** | phage terminase small subunit P27 family | UniRef100_A0A8E0DFY4 |
| **LdhD** | D-lactate dehydrogenase | UniRef100_A0A0D1I5H9 |
| **RocF** | arginase | UniRef100_UPI000C6D852F |
| **SasG** | E domain-containing protein | UniRef100_UPI000302884A |
| **SdrD** | MSCRAMM family adhesin SdrD | UniRef100_UPI00045EF767 |
| **Sph** | sphingomyelin phosphodiesterase | UniRef100_UPI0002C9D5E0 |

Proteins were considered consistently down-regulated if they were significantly down-regulated in 3 (out of 4) of the later timepoint isolate's secretome compared to T1. † proteins encoded by Sa2int prophage ‡ UniRef reference given in UniRef100 or UniRef100

**Table 3.**
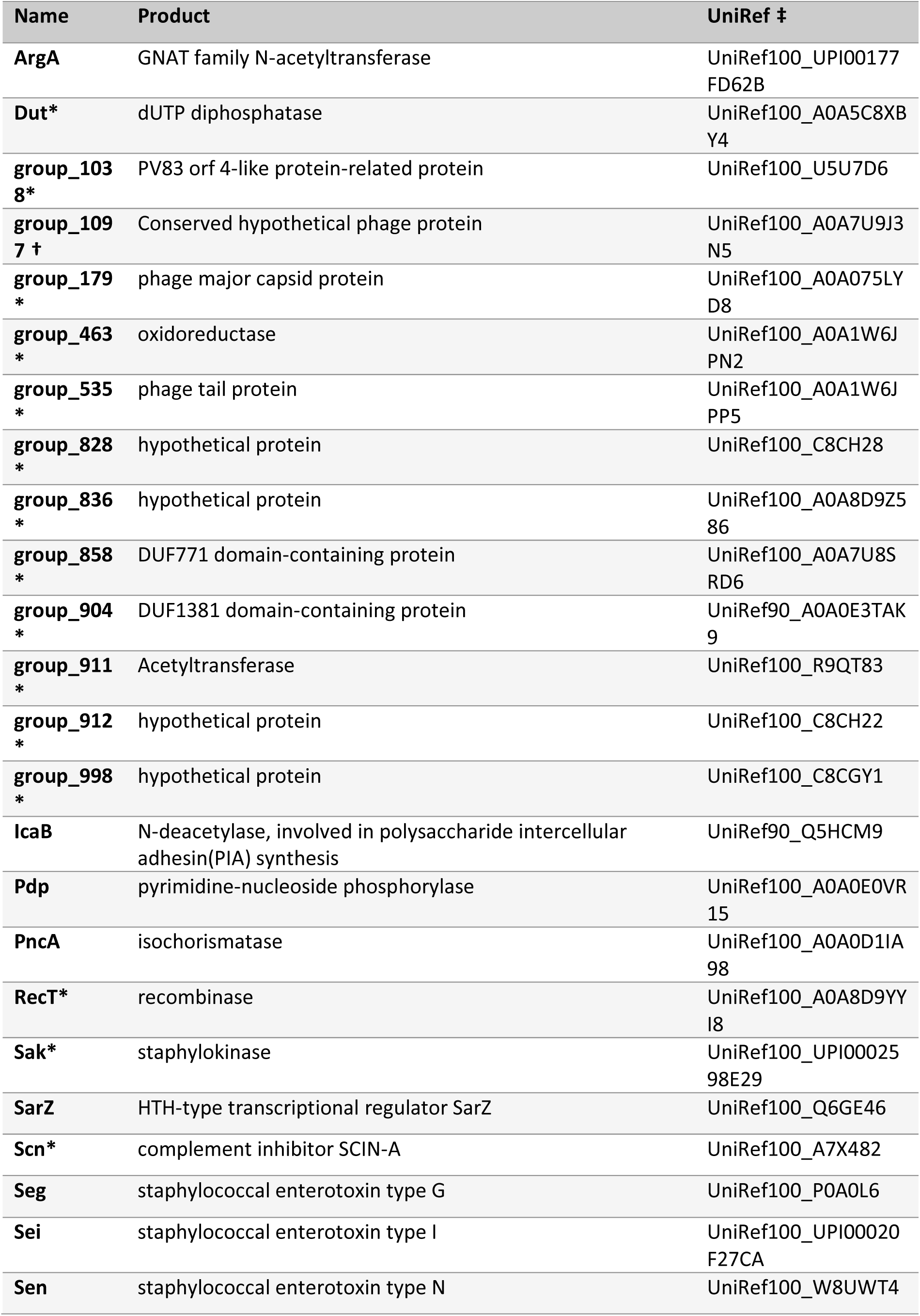
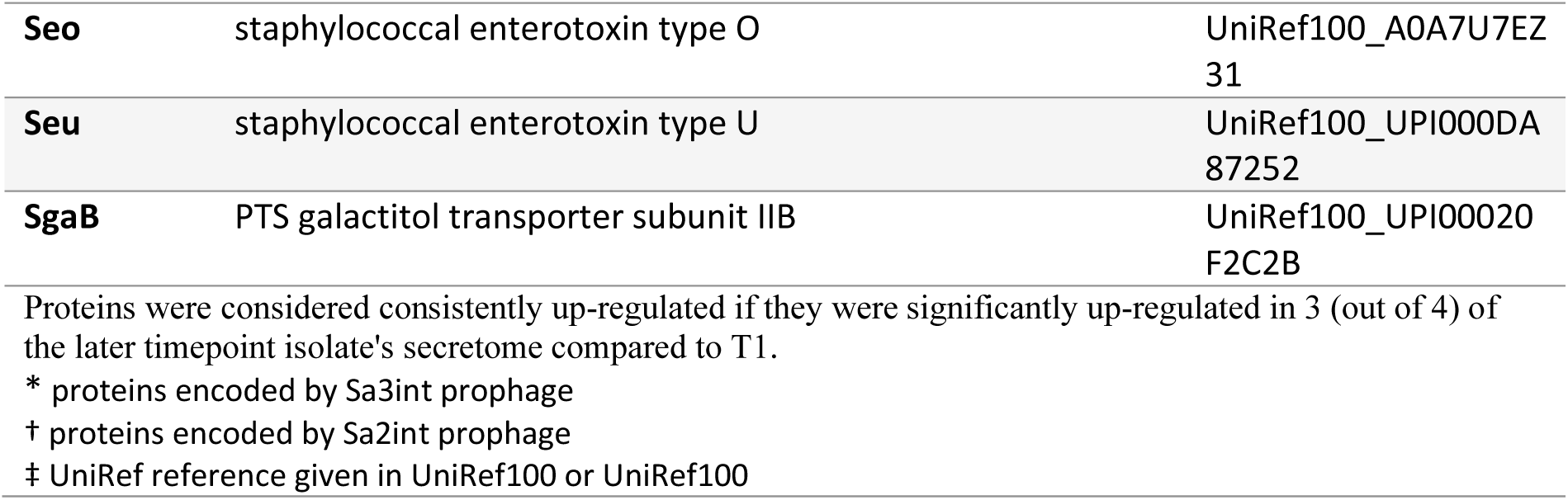
Consistent upregulation of proteins in the secretome of later timepoint isolates (T3, T5, T6, T8) compared to T1.

Out of the 26 consistently up-regulated proteins in the secretome of the later timepoint isolates, 15 were encoded on the genome of the Sa3int prophage. In addition, enterotoxin proteins such as Seg, Sei, Sen, Seo, and Seu were consistently upregulated, indicating an increased secretion of virulence factors by those isolates. Interestingly, the SarZ protein, a general promoter of virulence factors, was also consistently upregulated (Dyzenhaus et al., 2023; Tamber and Cheung, 2009). We also observed a consistent upregulation of the IcaB protein, which catalyses the N-deacetylation of PIA. N-deacetylation is required for bacterial cell surface retention of the PIA polymer (Vuong et al., 2004).

### Allelic exchange indicated mutations in *mfd* underlie a switch to hyper-biofilm forming mucoid phenotype

To determine the phenotypic consequences of the direct repeat sequence deletion in the *mfd* gene of T3 (Leu763fs), we employed allelic exchange to introduce this mutation into isolate T1, yielding isolate T1*^mfd^*^-T3^. To confirm the introduction of the mutation, we performed whole genome sequencing and read mapping with manual inspection which validated the presence of the intended deletion in the *mfd* gene of the T1 isolate (Supplementary Figure S6). The phenotypic impact was then assessed through examination of biofilm formation. The T1*^mfd^*^-T3^ mutant exhibited a significant increase in biofilm biomass to a similar level as isolate T3, when compared to T1 and RN4220 in TSB (p<0.05) (Figure 5A). Furthermore, when culturing the T1*^mfd^*^-T3^ mutant on congo red agar (CRA), we observed distinctive dark colonies, signifying the presence of PIA/PNAG, and a pronounced mucoid phenotype characterized by dry irregular edges, as shown in Figure 5B. Taken together, these experiments provide compelling evidence that the acquisition of the *mfd* mutation is essential in the phenotypic transformation towards a mucoid, hyper-biofilm formation in the longitudinal (T3-T8) clinical isolates.

**Figure 5.**
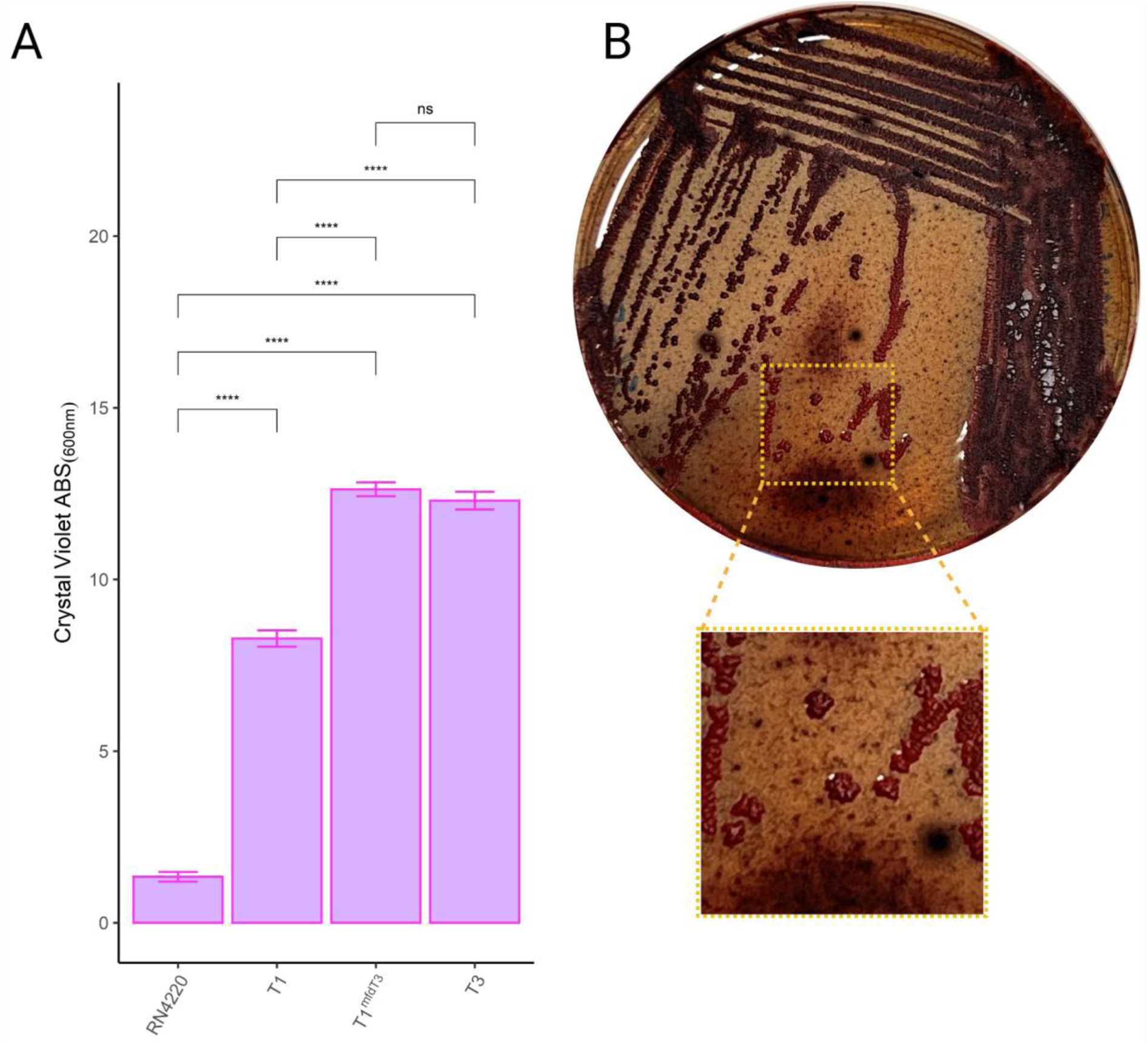
Impact of a clinical *mfd* mutation on biofilm and PIA production. (A) Biofilm quantification of two independently isolated T1*^mfd-^*^T3^ clone, T1, T3 and RN4220 using the crystal violet assay on TSB grown cells. (B) Congo red agar plate of T1*^mfd-^*^T3^ isolate, zoomed-in image of single colonies. Statistical significance was determined using one-way ANOVA and pairwise post-hoc T-test with Benjamini-Hochberg p-value adjustment (****p<0.0001; ns=non-significant).

To understand the impact of mutation combination, select mutations were introduced into the USA300 *S. aureus* strain JE2 (Table 4). None of the mutants exhibited a mucoid phenotype on CRA, ruling out the individual or combined roles of *icaR* and *mfd* genes in developing a mucoid phenotype in JE2. This suggests that the mucoid phenotype likely requires additional factors or genetic context present in T1 but not in JE2. Additionally, the deletion of either *icaR*, *mfd*, or the introduction of the Leu763fs into the *mfd* gene did not induce clumping in TSB, suggesting that hyper-biofilm formation is not regulated by these genes independently. However, the simultaneous deletion of *icaR* and *mfd* from JE2 led to increased clumping in TSB, resembling hyper-biofilm forming isolates, which highlights an interplay between *icaR* and *mfd* in the regulation of hyper-biofilm formation. Similarly, a mutant with a complete *icaR* deletion and an 8 bp deletion in *mfd* (JE2Δ*icaR mfd^Leu763fs^*) displayed comparable clumping, indicating that the truncation of *mfd* has an impact similar to the complete deletion (Figure 6). Further, antibiotic susceptibility to various antibiotics was similar across mutants and versus the parent isolate (Supplementary Table S4).

**Figure 6.**
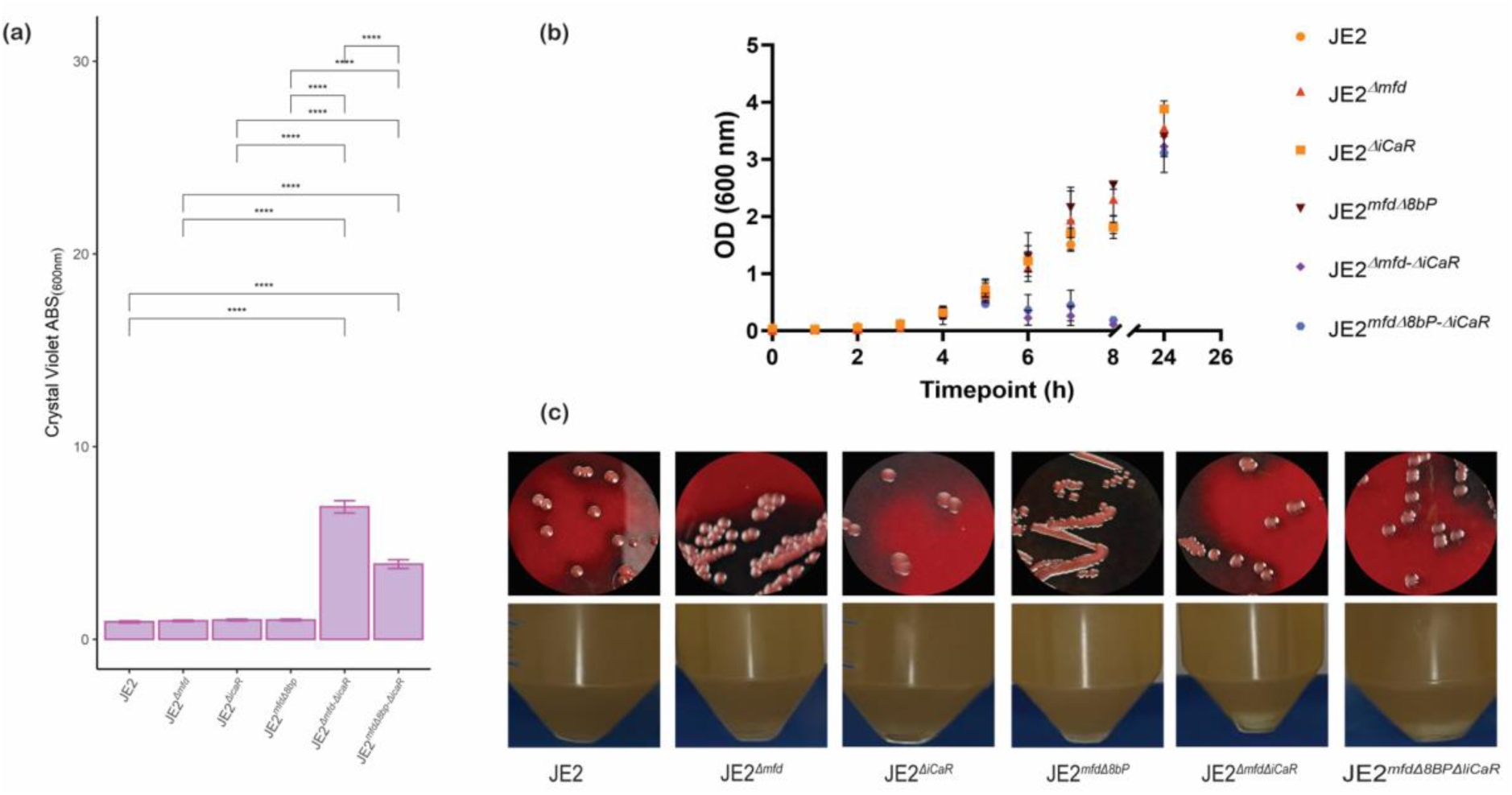
(a) Comparison of biofilm biomass of *S. aureus* mutants in the JE2 background. Single deletions of *mfd*, *icaR* and 8bp in *mfd* gene did not significantly change biofilm biomass compared to JE2, while combined deletions in both *mfd* and *icaR* genes resulted in significant rise in biofilm biomass. (b) growth curve of *S. aureus* mutants compared to JE2. combined deletions in both *mfd* and *icaR* genes resulted in bacterial clumping evidenced by reduction in OD readings after 5 h incubation followed by increase in readings after 24 h. (c) morphology of *S. aureus* mutant’s colonies on Congo red agar and in liquid culture. On Congo red agar all mutants displayed a non-mucoid phenotype regardless of mutation, while in liquid culture clumps were visible for mutants with deletions in both *mfd* and *icaR* genes. Statistical significance was determined using one-way ANOVA and pairwise post-hoc T-test with Benjamini-Hochberg p-value adjustment (****p<0.0001; ns=non-significant).

**Table 4.**
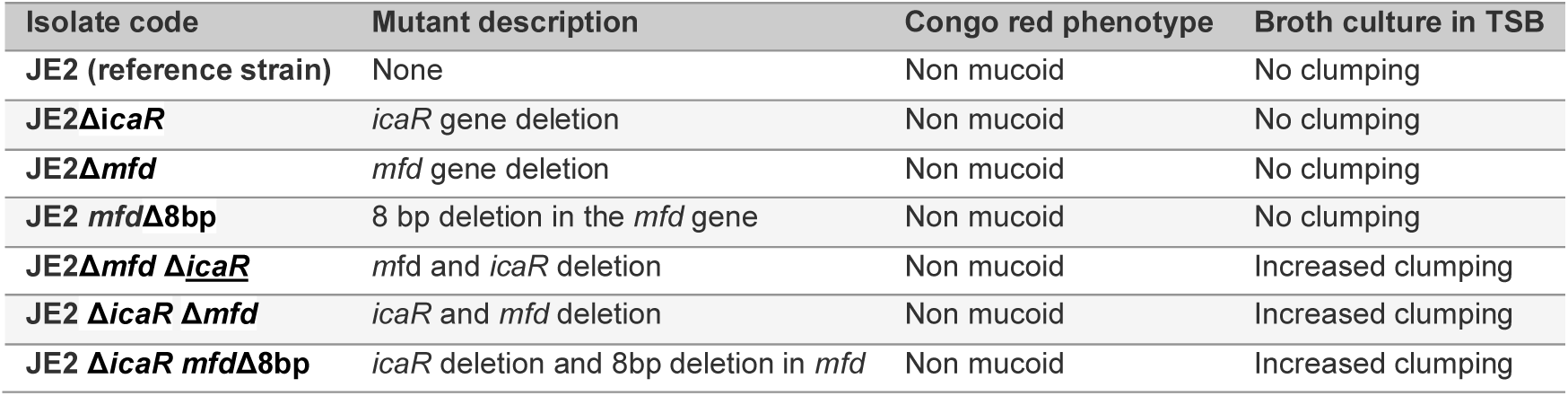

## Discussion

*S. aureus* is commonly isolated from the sinonasal cavity of CRS patients. The pathogen is likely to play an aggravating role in a subpopulation of CRS patients and other chronic inflammatory diseases such as atopic dermatitis and asthma (Teufelberger et al., 2019). Biofilm formation has been suspected of playing a pivotal role in the persistence of *S. aureus* and is an independent modifier of disease in CRS (Flemming *et al*., 2016; Psaltis et al., 2008). Here we show the within-host evolution of an *S. aureus* isolate yielding hyper-biofilm formation in a recalcitrant CRS patient.

Only a handful of studies have investigated the mucoid colony phenotype leading to increased biofilm production by *S. aureus*, with these variants occasionally isolated from the lungs of cystic fibrosis patients (Lennartz *et al*., 2019; Schwartbeck *et al*., 2016). To our knowledge, this is the first report of mucoid *S. aureus* isolates cultured from the sinonasal cavity of a CRS patient. In this study we utilised highly accurate closed genomes to analyse the relatedness of the longitudinal *S. aureus* clinical isolates. One non-mucoid isolate (T2) was not related to the other seven isolates, suggesting the introduction of a competing *S. aureus* strain in the niche at this timepoint. However, only mucoid *S. aureus* isolates closely related to T1 were isolated at time points subsequent to T2, indicating the dominance of T3-T8 over T2 and a potential fitness advantage of the mucoid, hyper biofilm-forming *S. aureus* compared to the non-mucoid *S. aureus* isolate. The survival advantage of the mucoid isolates was corroborated by the failure of multiple antibiotic courses to eradicate the pathogen.

Biofilm formation by *S. aureus* is a multi-staged process with many factors including genetic and environmental regulating the production (Hall-Stoodley et al., 2004; Raafat et al., 2019). The bacterial colonies expand by dividing and producing EPS, which enhance bacterial adhesion and gives rise to the biofilm matrix. The primary EPS of *S. aureus* biofilms is PNAG, which is regulated by the *icaR-icaADBC* locus. In line with other reports (Jefferson *et al*., 2003), we identified an inherited mutation (5’ IS21 insertion) inactivating the *icaR* repressor in the high biofilm forming T1, T3-T8 *S. aureus* isolates (Brooks and Jefferson, 2014; Jefferson *et al*., 2003; Schwartbeck *et al*., 2016).

However, disruption of the *icaR* by IS21 alone did not result in the mucoid *S. aureus* phenotype. Brooks and Jefferson reported difficulty maintaining pure mucoid cultures of their PNAG-overproducing laboratory strain *in vitro* as PNAG-negative phase mutants developed relatively quickly (Brooks and Jefferson, 2014). The authors proposed that this resulted from high-level production of PNAG which resulted in a fitness cost. PNAG-negative mutants contained a “TTTA” repeat insertion in the *icaC* gene, resulting in a premature stop codon (Brooks and Jefferson, 2014). Our results indicate that high-level production of PNAG might be advantageous *in vivo*, specifically in cavernous spaces of the body, such as the sinuses and lungs. However, overnight *in vitro* cultures of later timepoint mucoid isolates contained a mixture of aggregated and non-aggregated cells, indicating a potential phase shift. Furthermore, DNA sequence variation in the *icaC* gene through “TTTA” repeat insertion could be identified by short read mapping (T4, T7 and T8; Supplementary Figure 3) suggesting the presence of phase variation reversion mutants in the culture during DNA extraction.

Furthermore, our results showed that enhanced biofilm production was accompanied by the rise of autoaggregation of bacterial cells during the exponential growth phase. Therefore, the increased biofilm production is likely driven by the enhanced ability to aggregate. The initial stage of biofilm formation requires the aggregation of planktonic bacterial cells or their attachment to surfaces (biotic and abiotic). PNAG plays a pivotal role in mediating cell-cell adhesion and bacterial aggregation. However, little is known about the genetic factors that promote PIA/PNAG dependent aggregation. In a parallel project, we successfully transduced the Sa3int prophage found in later timepoints (T3-T8) into the T1 isolate. However, the lysogen-converted T1 isolate did not display the mucoid phenotype which ruled out any significant role for the prophage encoded genes (Nepal *et al*., 2025).

We successfully generated a T1 mutant harbouring the truncated *mfd* gene present in isolates T3-T8. The *mfd* gene encodes a conserved bacterial protein responsible for facilitating transcription-coupled DNA repair (Adebali et al., 2017). Strikingly, these mutants exhibited phenotypic characteristics closely resembling the T3-T8 mucoid isolates both in terms of biofilm formation and colony morphology. Our analysis highlights the involvement of the *mfd* gene in modulating the mucoid phenotype. Interestingly, the clinical isolates T3-T8 carrying the truncated *mfd* genes persisted in the paranasal sinuses of the patient for an extended period. They even outcompeted the T1 isolate containing the full length *mfd* gene, even though Mfd induces resistance to the eukaryotic nitrogen response produced by macrophages (Darrigo et al., 2016). Previous research has demonstrated that reduced *mfd* expression in *S. aureus* resulted in a partial decrease in PNAG reactivity implying that the Mfd protein plays a role in modulating biofilm development (Tu Quoc et al., 2007). Considering the diverse functions of the Mfd protein, it is plausible that Mfd indirectly influences biofilm formation by regulating the expression of genes associated with this process. Alterations in gene expression profiles may have an impact on the production of biofilm-related components, including EPS and adhesins. We recapitulated these results in the JE2 background that did not exhibit a mucoid phenotype. Sequential acquisition of mutations in *icaR* and *mfd*, or the combination of the 8 bp deletion in *mfd* with *icaR* deletion, led to increased clumping in TSB and enhanced biofilm production. These findings reaffirm the established role of *icaR* in biofilm formation and indicate a functional interaction between *icaR* and *mfd*. Nevertheless, in the genetic context of JE2 we were unable to fully induce a mucoid phenotype present in isolates T3-T8.

The specific mechanisms connecting the Mfd protein to biofilm formation through the formation of mucoid *S. aureus* isolates is an ongoing subject of research, and further studies are required to gain a comprehensive understanding of this relationship. Moreover, the association between Mfd and biofilm formation may vary among different *S. aureus* strains, as the regulatory networks involved can be intricate and underexplored.

Cell-surface-associated adhesins facilitate the attachment of cells to their surroundings (Koo et al., 2017; Sauer et al., 2022). For *S. aureus*, these are mainly the ‘microbial surface components recognising adhesive matrix molecules’ (MSCRAMMs) family proteins such as the fibronectin binding proteins (FnBP) and SasG, (Foster et al., 2014). Our secretome data showed a reduced presence of cell wall associated (CWA) proteins, including SasG, FnbA and SdrD. As CWA proteins have been reported to be released from the cell wall during cell growth, this could indicate increased retention of CWA proteins (Becker et al., 2014). Alternatively, there could be an overall down-regulation of CWA proteins in later timepoint isolates (T3, T5, T6, T8) vs T1, in line with the consistent up-regulation of virulence factors (Seg, Sei, Sen, Seo, Seu) and SarZ (Arciola *et al*., 2015; Tamber and Cheung, 2009). Interestingly, SarZ, a major regulator of virulence factors, was consistently identified and found to be overexpressed in the secretome of later timepoint isolates vs T1. This is surprising as SarZ is a DNA-binding protein not known to be secreted by *S. aureus* (Arciola *et al*., 2015; Tamber and Cheung, 2009). These findings might be indicative of an overall overexpression of SarZ in those isolates. Further experiments are required to test this hypothesis.

We also found a reduced secretion of proteins encoded by the Sa2int prophage in the mucoid isolates. The underlying mechanism for this observation is currently unknown, however, it may be associated with the lysogenisation of the later-timepoint mucoid isolates by the Sa3int prophage, leading to polylysogeny and potential selection between phages within the bacterial host (Burns et al., 2015). Polylysogeny can result in discrete cell subsets producing either individual (or more rarely a combination) of prophages triggered by certain stimuli, suggesting that the acquisition of the Sa3int prophage reduces the proportion of cells producing active Sa2int compared to the T1 isolate (Silpe et al., 2023; Wang et al., 2023). While the secretion of Sa3int prophage-carrying virulence factors *scn*, *chp* and *sak* at later timepoints is not surprising as Sa3int prophages are known to be critical in human colonisation (Chaguza *et al*., 2022), the maintenance of the Sa2int prophage over time suggests that Sa2int must either confer a survival advantage, or that there may be certain stimuli present in the sinonasal niche that only triggers lysis in Sa2int as compared to Sa3int. Further studies are necessary to fully understand the impact of prophage integration on protein secretion by *S. aureus* and its relevance to chronic rhinosinusitis. Overall, these results add to our understanding of the complex interplay between prophages and bacterial evolution in chronic infections.

## Conclusion

Our results show the genetic within-host evolution towards a mucoid *S. aureus* phenotype. Over time, the isolate evolved into a mucoid, hyper-biofilm former with increased secretion of virulence factors, possibly conferring survival fitness to the bacteria within the host. Allelic exchange experiments confirmed that mutation of the *mfd* gene could activate the mucoid phenotype.

---

**Figure S1.**
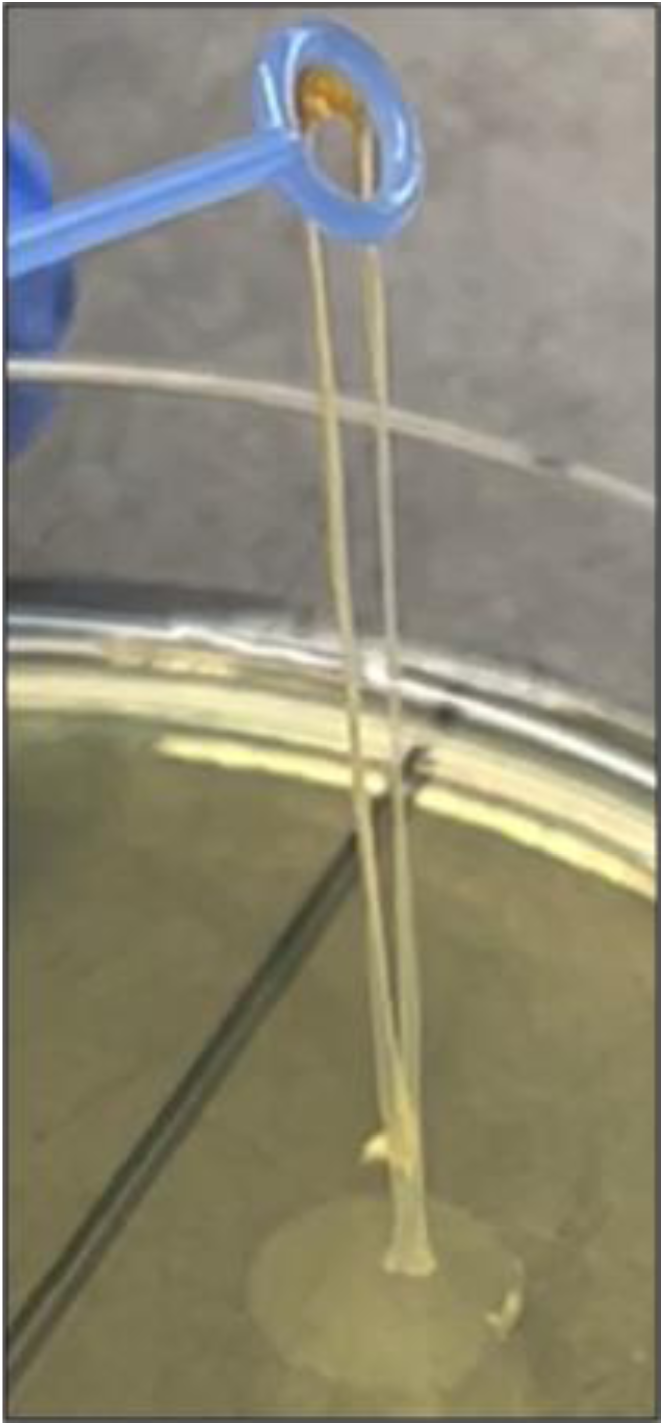
Mucoid *S. aureus* (T3) grown overnight on tryptic soy agar; the adhesion of bacterial plaque is shown by touching bacterial plaque with a loop.

**Figure S2.**
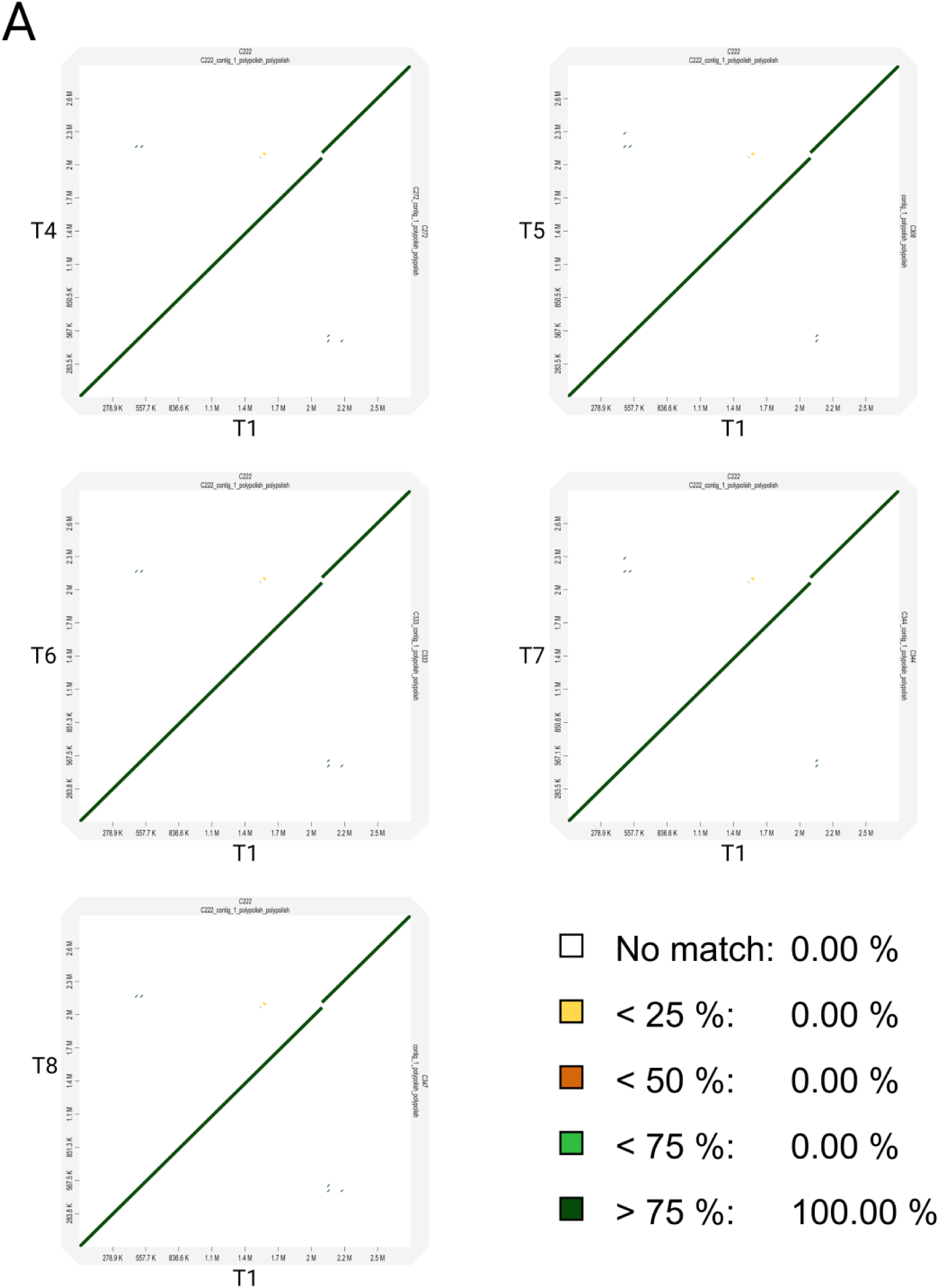
Pairwise genome alignment of clinical isolates. (A) Dot plot comparing T1 to T4-T8. The identity percentage of the BLAST-like alignment is depicted using colour coding.

**Figure S3.**
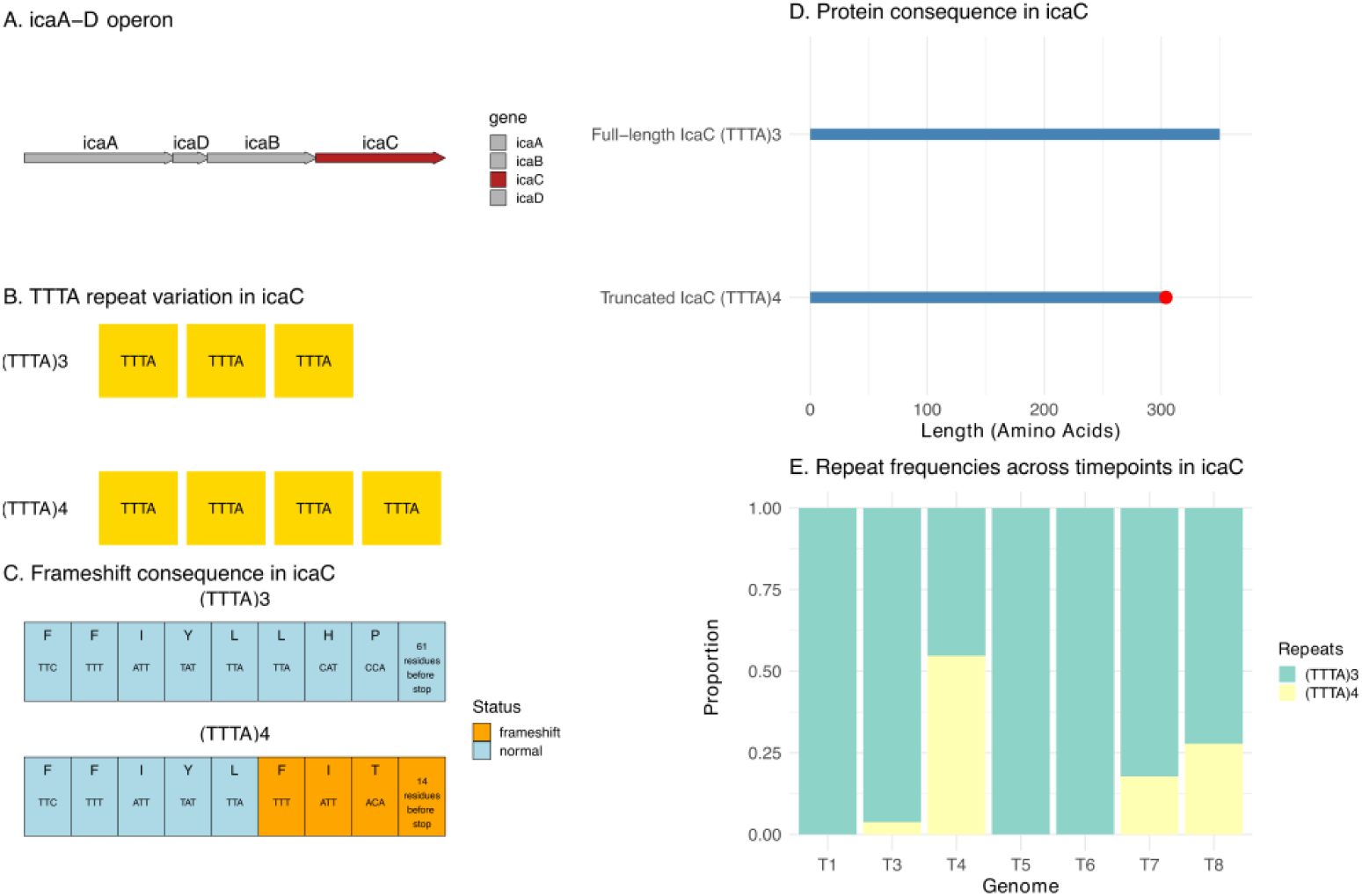
TTTA repeat variation in *icaC*. (A) Schematic map of *icaA-C* locus. (B) TTTA repeat variation in *icaC* between three i.e. (TTTA)3 and four i.e. (TTTA)4 copies. (C) Consequence of (TTTA)4 compared to (TTTA)3 on the residues (top) and nucleotides (bottom, in codon triplets) of the fourth TTTA repeat. All positions following the frameshift are indicated in yellow tiles. (D) Length truncation of (TTTA)3 (350 amino acids) compared to (TTTA)4 (304 amino acids, truncation indicated in red for effect). (E) Proportions of (TTTA)3 and (TTTA)4 in short reads across timepoints from T1 to T8. Notably, T3 (3.7%), T4 (54.5%), T7 (17.8%) and T8 (27.8%) had subpopulations of (TTTA)4.

**Figure S4.**
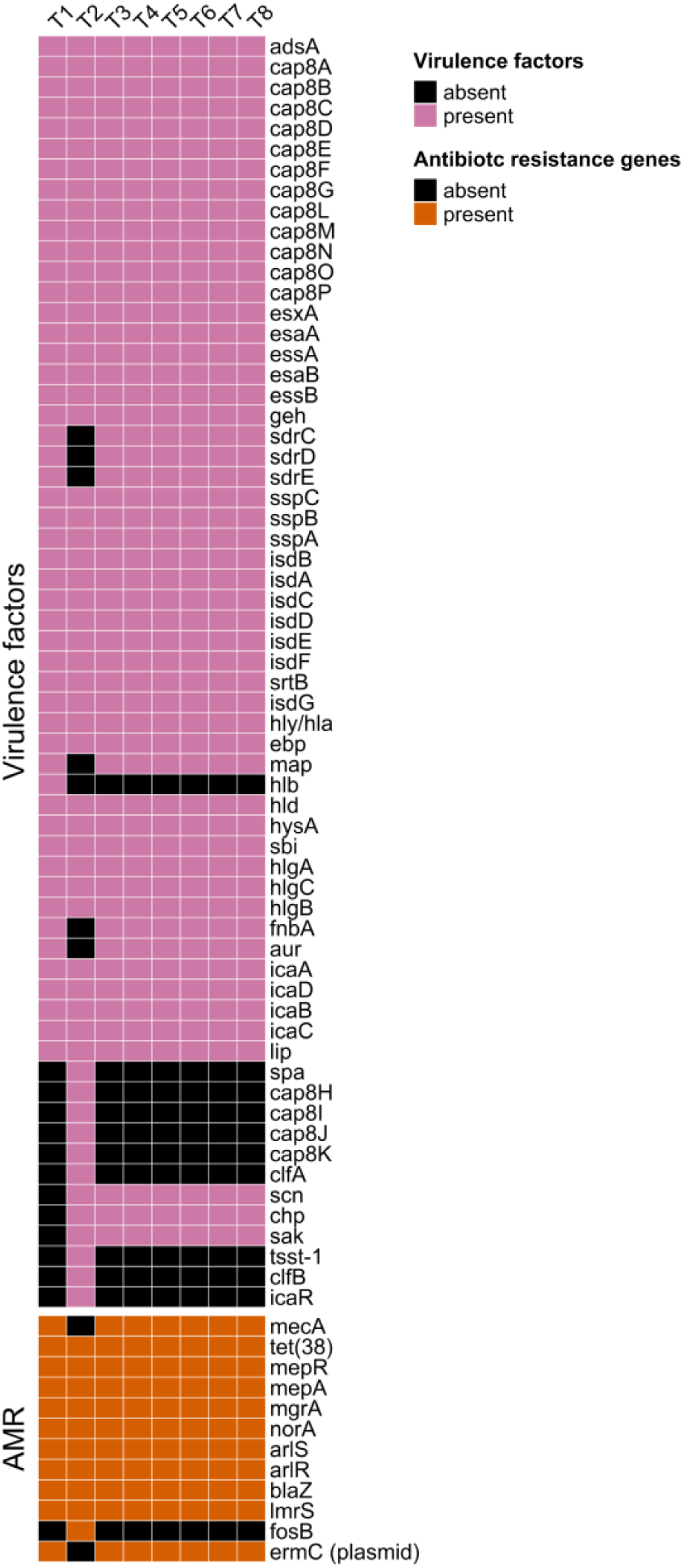
Virulence factors and antimicrobial resistance genes detected in the genome of the isolates.

**Figure S5.**
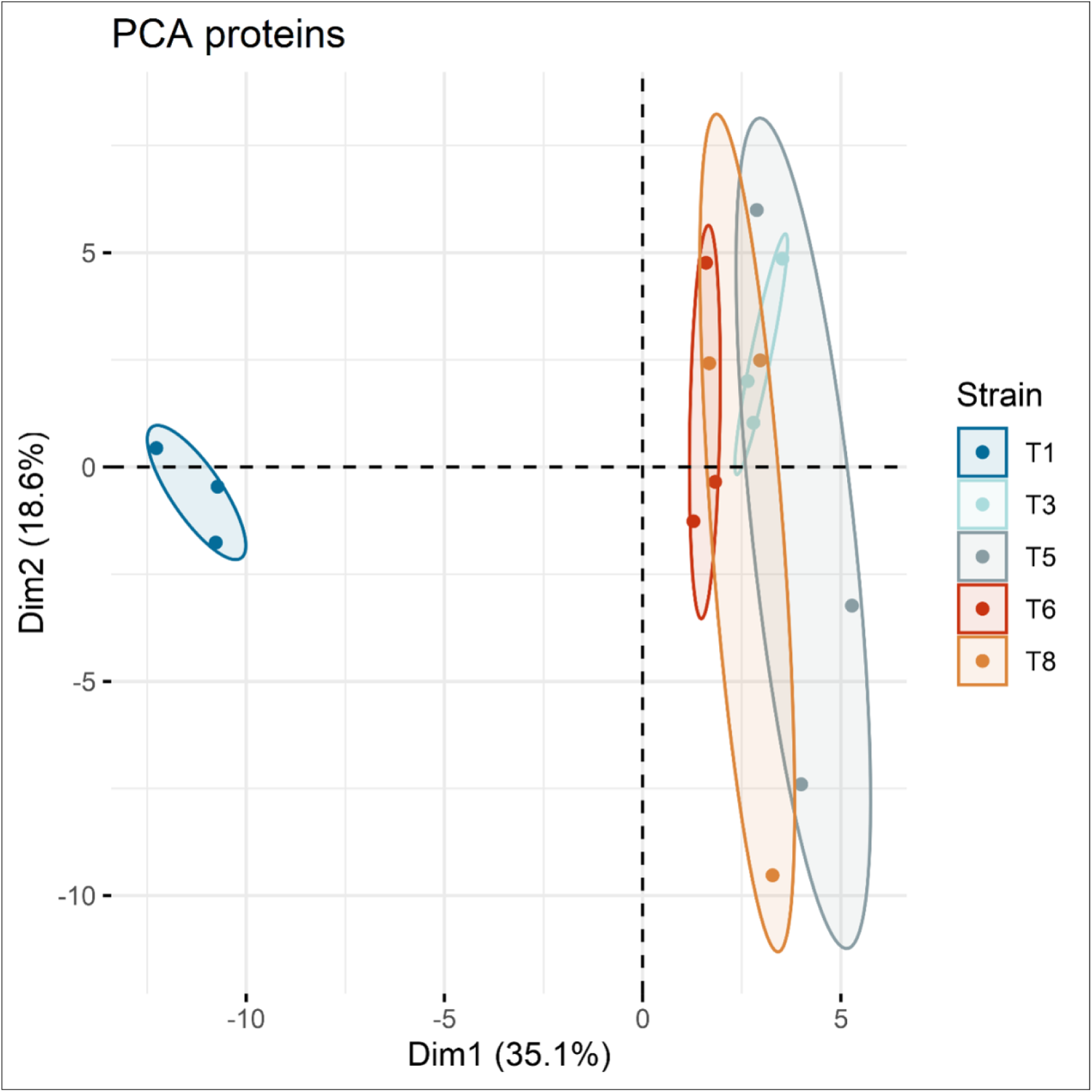
Principal component analysis (PCA) of mucoid *S. aureus* isolates. The PCA is based on the top 100 most variable proteins, and each point represents a sample colour-coded by group. The numbers in parentheses on the x- and y-axis represent the percentage of variation explained by PC1 and PC2, respectively.

**Figure S6.**
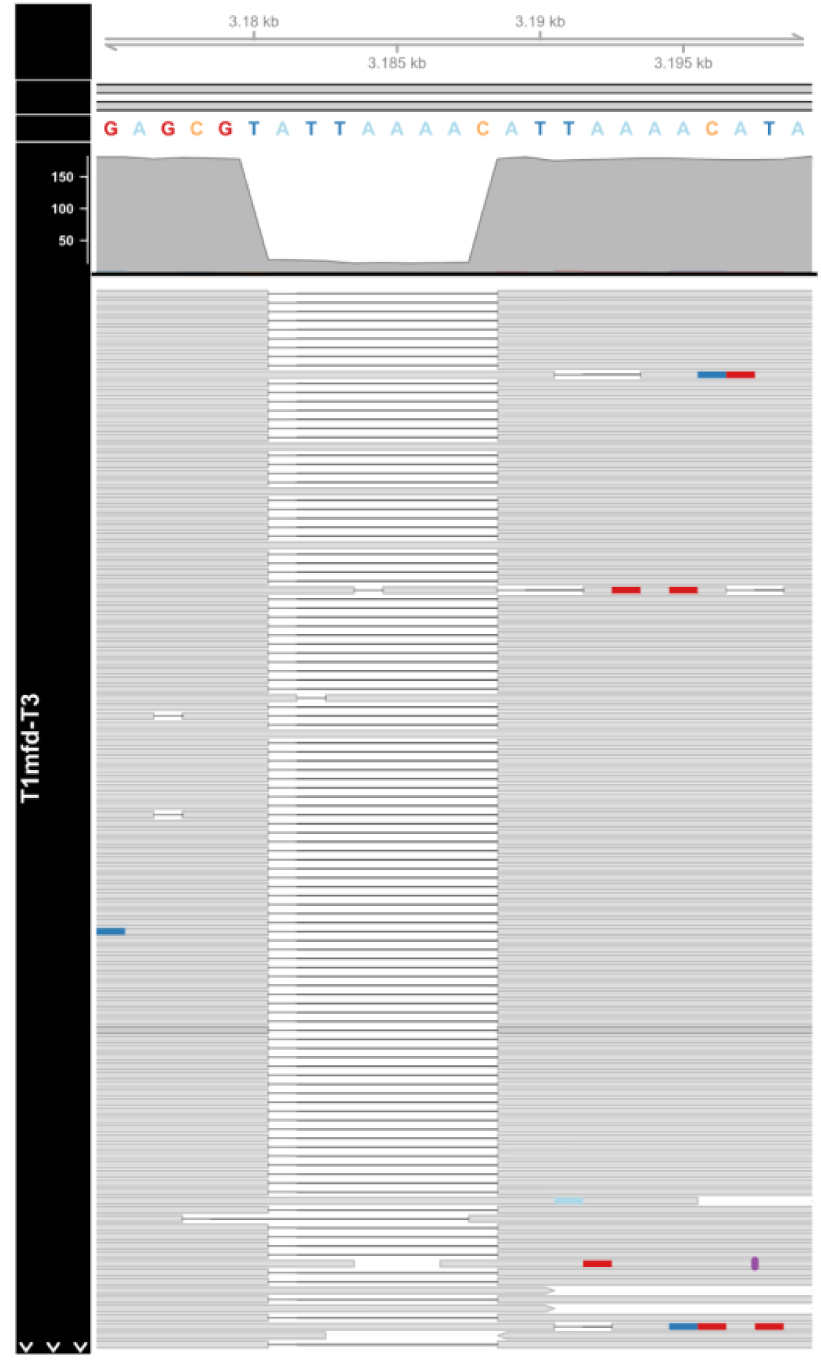
Visualization of long reads mapping to the *mfd* gene in isolate T1 *^mfd^*^-T3^. Pile-up plots of long reads from T1 *^mfd^*^-T3^ mapped T1 reference isolate. The 8bp deletion can be observed in T1 *^mfd^*^-T3^ isolate sequenced.

**Figure S7.**
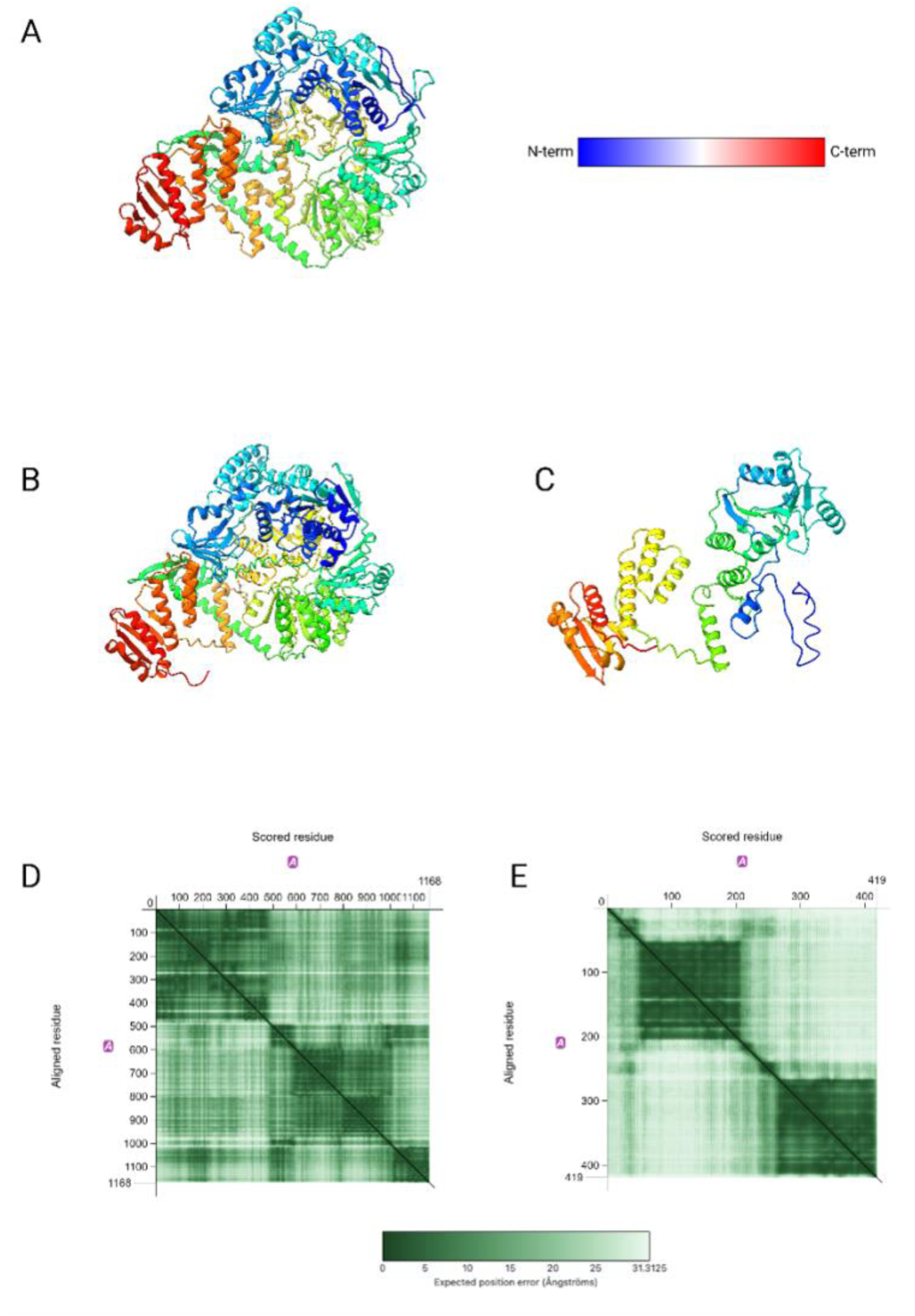
(A) Top-view depiction presenting the α-carbon backbone ribbon model of the *E. coli* Mfd protein’s X-ray crystal structure, as experimentally determined at 3.2 Å resolution (Deaconescu and Darst, 2005). The color gradation is from blue N-terminus to red C-terminus. (B) Colabfold-predicted structure of the T1 Mfd protein. (C) Colabfold-predicted structure of the T3 Mfd protein. (D) Predicted Aligned Error of the T1 Mfd protein. Green denotes lower error, white signifies areas of uncertainty. (E) Predicted Aligned Error of the T3 Mfd protein, revealing the estimated accuracy of multimer folding. Green represents lower error, white indicates regions of uncertainty.

## Acknowledgements

We thank members of The Department of Surgery-Otolaryngology Head and Neck Surgery at Adelaide University for their continuous support throughout this project. GH is supported by Adelaide University and The Hospital Research Foundation scholarships. The work was funded through a Passe and Williams Foundation Senior Fellowship to SV and an NHMRC investigator grant (APP1196832) to PJW.

## Competing interests

The authors declare no competing interests that are relevant to this work.

## Abbreviations

CC: clonal complex
CI: clinical isolate
CRA: Congo Red Agar
CRS: chronic rhinosinusitis
CRSwNP: chronic rhinosinusitis with nasal polyps
CV: crystal violet
CWA: cell-wall anchored
DEP: differentially expressed proteins
DIA-MS: Data-independent acquisition mass spectrometry
EPS: extracellular polymeric substance
gDNA: genomic DNA
LC-MS: Liquid Chromatography-Mass Spectrometry
MFU: McFarland standard unit
MGEs: mobile genetic elements
MRSA: methicillin-resistant *Staphylococcus aureus*
MSCRAMMs: microbial surface components recognizing adhesive matrix molecules
NA: nutrient agar
OD_600_: optical density at 600 nm
PIA: polysaccharide intercellular adhesin
PNAG: poly-β(1-6)-N-acetylglucosamine
SNP: single nucleotide polymorphism
tRNA: transfer RNA

---

**Table S1.** Crystal violet assay results.

| Clinical isolates | Mean absorbance (OD<sub>600</sub>) | Standard error | Ratio (CI/RN4220) | Growth media |
| --- | --- | --- | --- | --- |
| T1 | 1.92 | 0.11 | 1.95 | MHB |
| T2 | 0.3 | 0.03 | 0.3 | MHB |
| T3 | 4.93 | 0.4 | 5 | MHB |
| T4 | 4.67 | 0.19 | 4.73 | MHB |
| T5 | 4.32 | 0.28 | 4.38 | MHB |
| T6 | 3.91 | 0.45 | 3.96 | MHB |
| T7 | 4.18 | 0.15 | 4.24 | MHB |
| T8 | 4.78 | 0.18 | 4.85 | MHB |
| RN4220 | 0.99 | 0.1 | 1 | MHB |
| T1 | 7.07 | 0.24 | 1.87 | TSB |
| T2 | 1.29 | 0.04 | 0.34 | TSB |
| T3 | 10.97 | 0.06 | 2.9 | TSB |
| T4 | 10.98 | 0.34 | 2.9 | TSB |
| T5 | 10.81 | 0.32 | 2.85 | TSB |
| T6 | 11.04 | 0.49 | 2.91 | TSB |
| T7 | 10.95 | 0.39 | 2.89 | TSB |
| T8 | 11 | 0.22 | 2.9 | TSB |
| RN4220 | 3.79 | 0.66 | 1 | TSB |
| T1 | 2.32 | 0.13 | 3.87 | NB |
| T2 | 0.14 | 0.04 | 0.24 | NB |
| T3 | 3.36 | 0.22 | 5.62 | NB |
| T4 | 3.13 | 0.24 | 5.23 | NB |
| T5 | 3.4 | 0.32 | 5.67 | NB |
| T6 | 3.59 | 0.36 | 5.99 | NB |
| T7 | 3.48 | 0.36 | 5.82 | NB |
| T8 | 3.29 | 0.12 | 5.5 | NB |
| RN4220 | 0.6 | 0.1 | 1 | NB |

Note: MHB, Muller-Hinton broth; TSB, Tryptic soy broth; NB, Nutrient broth; n=3 replicates

**Table S2.** Structural evolutionary changes between T1 and later timepoint isolates (T3-T8)

| Reference isolate | Start | End | Name | length | Query isolate | Coordinate | Repeated region |
| --- | --- | --- | --- | --- | --- | --- | --- |
| **T1(C222)** | 1365848 | 1365848 | insertion | 2197 | T3(C247) | 1365841-1368037 | NA |
| **T1(C222)** | 1365848 | 1365848 | duplication | 6 | T3(C247) | 1368038-1368043 | 1365843-1365848 |
| **T1(C222)** | 1943289 | 1943289 | insertion | 227 | T3(C247) | 1945485-1945711 | NA |
| **T1(C222)** | 1943289 | 1943289 | duplication | 77 | T3(C247) | 1945712-1945788 | 1943213-1943289 |
| **T1(C222)** | 2042009 | 2042009 | insertion | 43793 | T3(C247) | 2044511-2088303 | NA |
| **T1(C222)** | 2042009 | 2042009 | duplication | 14 | T3(C247) | 2088304-2088317 | 2041996-2042009 |
| **T1(C222)** | 1365848 | 1365848 | insertion | 2197 | T4(C272) | 1365806-1368002 | NA |
| **T1(C222)** | 1365848 | 1365848 | duplication | 6 | T4(C272) | 1368003-1368008 | 1365843-1365848 |
| **T1(C222)** | 1943289 | 1943289 | insertion | 227 | T4(C272) | 1945450-1945676 | NA |
| **T1(C222)** | 1943289 | 1943289 | duplication | 77 | T4(C272) | 1945677-1945753 | 1943213-1943289 |
| **T1(C222)** | 2042009 | 2042009 | insertion | 43793 | T4(C272) | 2044474-2088266 | NA |
| **T1(C222)** | 2042009 | 2042009 | duplication | 14 | T4(C272) | 2088267-2088280 | 2041996-2042009 |
| **T1(C222)** | 1365848 | 1365848 | insertion | 2197 | T5(C308) | 1365842-1368038 | NA |
| **T1(C222)** | 1365848 | 1365848 | duplication | 6 | T5(C308) | 1368039-1368044 | 1365843-1365848 |
| **T1(C222)** | 1943289 | 1943289 | insertion | 227 | T5(C308) | 1945486-1945712 | NA |
| **T1(C222)** | 1943289 | 1943289 | duplication | 77 | T5(C308) | 1945713-1945789 | 1943213-1943289 |
| **T1(C222)** | 2042009 | 2042009 | insertion | 43793 | T5(C308) | 2044510-2088302 | NA |
| **T1(C222)** | 2042009 | 2042009 | duplication | 14 | T5(C308) | 2088303-2088316 | 2041996-2042009 |
| **T1(C222)** | 610668 | 610668 | insertion | 740 | T6(C333) | 610661-611400 | NA |
| **T1(C222)** | 610668 | 610668 | duplication | 3 | T6(C333) | 611401-611403 | 610666-610668 |
| **T1(C222)** | 1365848 | 1365848 | insertion | 2197 | T6(C333) | 1366585-1368781 | NA |
| **T1(C222)** | 1365848 | 1365848 | duplication | 6 | T6(C333) | 1368782-1368787 | 1365843-1365848 |
| **T1(C222)** | 1567348 | 1567348 | insertion | 2197 | T6(C333) | 1570288-1572484 | NA |
| **T1(C222)** | 1567348 | 1567348 | duplication | 6 | T6(C333) | 1572485-1572490 | 1567343-1567348 |
| **T1(C222)** | 1943289 | 1943289 | insertion | 227 | T6(C333) | 1948432-1948658 | NA |
| **T1(C222)** | 1943289 | 1943289 | duplication | 77 | T6(C333) | 1948659-1948735 | 1943213-1943289 |
| **T1(C222)** | 2042009 | 2042009 | insertion | 43793 | T6(C333) | 2047456-2091248 | NA |
| **T1(C222)** | 2042009 | 2042009 | duplication | 14 | T6(C333) | 2091249-2091262 | 2041996-2042009 |
| **T1(C222)** | 2362030 | 2362328 | collapsed_tandem_repeat | NA | T6(C333) | 2411282 | NA |
| **T1(C222)** | 610668 | 610668 | insertion | 739 | T7(C344) | 610661-611399 | NA |
| **T1(C222)** | 610668 | 610668 | duplication | 3 | T7(C344) | 611400-611402 | 610666-610668 |
| **T1(C222)** | 1365848 | 1365848 | insertion | 2197 | T7(C344) | 1366584-1368780 | NA |
| **T1(C222)** | 1365848 | 1365848 | duplication | 6 | T7(C344) | 1368781-1368786 | 1365843-1365848 |
| **T1(C222)** | 1943289 | 1943289 | insertion | 227 | T7(C344) | 1946228-1946454 | NA |
| **T1(C222)** | 1943289 | 1943289 | duplication | 77 | T7(C344) | 1946455-1946531 | 1943213-1943289 |
| **T1(C222)** | 2042009 | 2042009 | insertion | 43793 | T7(C344) | 2045252-2089044 | NA |
| **T1(C222)** | 2042009 | 2042009 | duplication | 14 | T7(C344) | 2089045-2089058 | 2041996-2042009 |
| **T1(C222)** | 2362030 | 2362328 | collapsed_tandem_repeat | NA | T7(C344) | 2409078 | NA |
| **T1(C222)** | 610668 | 610668 | insertion | 739 | T8(C347) | 610661-611399 | NA |
| **T1(C222)** | 610668 | 610668 | duplication | 3 | T8(C347) | 611400-611402 | 610666-610668 |
| **T1(C222)** | 1365848 | 1365848 | insertion | 2197 | T8(C347) | 1366584-1368780 | NA |
| **T1(C222)** | 1365848 | 1365848 | duplication | 6 | T8(C347) | 1368781-1368786 | 1365843-1365848 |
| **T1(C222)** | 1567348 | 1567348 | insertion | 2197 | T8(C347) | 1570287-1572483 | NA |
| **T1(C222)** | 1567348 | 1567348 | duplication | 6 | T8(C347) | 1572484-1572489 | 1567343-1567348 |
| **T1(C222)** | 1943289 | 1943289 | insertion | 227 | T8(C347) | 1948432-1948658 | NA |
| **T1(C222)** | 1943289 | 1943289 | duplication | 77 | T8(C347) | 1948659-1948735 | 1943213-1943289 |
| **T1(C222)** | 2042009 | 2042009 | insertion | 43793 | T8(C347) | 2047458-2091250 | NA |
| **T1(C222)** | 2042009 | 2042009 | duplication | 14 | T8(C347) | 2091251-2091264 | 2041996-2042009 |
| **T1(C222)** | 2362030 | 2362328 | collapsed_<br>tandem_re<br>peat | NA | T8(C347) | 2411284 | NA |

**Table S3.** The *S. aureus* mutational divergence between T1 and later timepoint isolates (T3-T8)

| Clinical isolates | Position | Mutation type | Reference | Strand | Gene | Gene product |
| --- | --- | --- | --- | --- | --- | --- |
| **T3(C247)** | 503584 | frameshift deletion | ATTAAAAC | + | *mfd* | transcription-repair coupling factor |
| **T3(C247)** | 1072853 | SNP | G | - | *isdA* | iron-regulated surface determinant protein A |
| **T3(C247)** | 1254003 | SNP | C | + | *ymdB* | metallophosphoesterase |
| **T4(C272)** | 503584 | frameshift deletion | ATTAAAAC | + | *mfd* | transcription-repair coupling factor |
| **T4(C272)** | 577808 | complex | CTCAG | + | *sdrC* | Ser-Asp rich fibrinogen-binding bone sialoprotein-binding protein |
| **T4(C272)** | 1072853 | SNP | G | - | *isdA* | iron-regulated surface determinant protein A |
| **T4(C272)** | 1254003 | SNP | C | + | *ymdB* | metallophosphoesterase |
| **T5(C308)** | 503584 | frameshift deletion | ATTAAAAC | + | *mfd* | transcription-repair coupling factor |
| **T5(C308)** | 1072853 | SNP | G | - | *isdA* | iron-regulated surface determinant protein A |
| **T5(C308)** | 1254003 | SNP | C | + | *ymdB* | metallophosphoesterase |
| **T6(C333)** | 503584 | frameshift deletion | ATTAAAAC | + | *mfd* | transcription-repair coupling factor |
| **T6(C333)** | 658592 | SNP | A | + | *tagX* | Putative glycosyltransferase TagX |
| **T6(C333)** | 1072853 | SNP | G | - | *isdA* | iron-regulated surface determinant protein A |
| **T6(C333)** | 1254003 | SNP | C | + | *ymdB* | metallophosphoesterase |
| **T6(C333)** | 2361859 | SNP | A | NA | NA | NA |
| **T6(C333)** | 2609421 | SNP | G | NA | NA | NA |
| **T7(C344)** | 503584 | frameshift deletion | ATTAAAAC | + | *mfd* | transcription-repair coupling factor |
| **T7(C344)** | 658592 | SNP | A | + | *tagX* | Putative glycosyltransferase TagX |
| **T7(C344)** | 1072853 | SNP | G | - | *isdA* | iron-regulated surface determinant protein A |
| **T7(C344)** | 1254003 | SNP | C | + | *ymdB* | metallophosphoesterase |
| **T7(C344)** | 2361859 | SNP | A | NA | NA | NA |
| **T7(C344)** | 2609421 | SNP | G | NA | NA | NA |
| **T8(C347)** | 503584 | frameshift deletion | ATTAAAAC | + | *mfd* | transcription-repair coupling factor |
| **T8(C347)** | 658592 | SNP | A | + | *tagX* | Putative glycosyltransferase TagX |
| **T8(C347)** | 1072853 | SNP | G | - | *isdA* | iron-regulated surface<br>determinant protein A |
| **T8(C347)** | 1254003 | SNP | C | + | *ymdB* | metallophosphoesterase |
| **T8(C347)** | 2361859 | SNP | A | NA | NA | NA |
| **T8(C347)** | 2609421 | SNP | G | NA | NA | NA |

Note: T1 is used as the reference isolate; SNP, Single nucleotide polymorphism

**Table S4.** Antibiotic sensitivity of different *S. aureus* isolates at different time points and *S. aureus* mutants compared to laboratory strain JE2 reported as minimum inhibitory concentration (MIC), R= resistant, S= susceptible (n = 3).

| Antibiotics | Amoxycillin/clavulanic<br>acid<br>mg/L | Vancomycin<br>mg/L | Mupirocin<br>mg/L | Levofloxacin<br>mg/L |
| --- | --- | --- | --- | --- |
| **T1 (C222)** | >64/16 (R) | 0.5 (S) | 0.125 (S) | 4 (R) |
| **T2 (C229)** | 32/8 (R) | 0.5 (S) | 0.125 (S) | 0.25 (S) |
| **T3 (C247)** | >64/16 (R) | 0.5 (S) | 0.125 (S) | 4 (R) |
| **T4 (C272)** | >64/16 (R) | 0.5 (S) | 0.125 (S) | 4 (R) |
| **T5 (C308)** | >64/16 (R) | 0.5 (S) | 0.125 (S) | 4 (R) |
| **T6 (C333)** | >64/16 (R) | 0.5 (S) | 0.125 (S) | 4 (R) |
| **T7 (C344)** | >64/16 (R) | 0.5 (S) | 0.125 (S) | 4 (R) |
| **T8 (347)** | >64/16 (R) | 0.5 (S) | 0.125 (S) | 4 (R) |
| **JE2** | 16/4 (R) | 1 (S) | 0.5 (S) | 8 (R) |
| **JE2 $\Delta mfd$** | 16/4 (R) | 1 (S) | 0.5 (S) | 8 (R) |
| **JE2 $\Delta icaR$** | 16/4 (R) | 1 (S) | 0.5 (S) | 8 (R) |
| **JE2 $\Delta mfd \Delta icaR$** | 16/4 (R) | 1 (S) | 0.5 (S) | 8 (R) |
| **JE2 $\Delta icaR \Delta mfd$** | 16/4 (R) | 1 (S) | 0.5 (S) | 8 (R) |
| **JE2 $mfd \Delta 8bp$** | 16/4 (R) | 1 (S) | 0.5 (S) | 8 (R) |
| **JE2 $mfd \Delta 8bp \Delta icaR$** | 16/4 (R) | 1 (S) | 0.5 (S) | 8 (R) |

## Supplementary texts

### ST1 Extraction of *S. aureus* genomic DNA

Briefly, 700 µL of overnight broth culture in TSB was centrifuged (4000 x g) in a microcentrifuge tube for 10 minutes. The pellet was suspended in 180 µL of enzymatic filter sterilised lysis buffer (20 mM Tris-Cl, pH8; 2mM sodium EDTA; 1.2% Triton X-100, 200 µg/ml final concentration lysostaphin) and incubated at 37 °C for 30 minutes. After that, samples were centrifuged with the DNeasy Mini spin column. Finally, the sample was eluted and stored at -20°C till use.

### ST2 Secretome digestion

The proteins were reduced with tris(2-carboxyethyl)phosphine (TCEP, 10 mM, 30 min, 56°C) and alkylated in the dark with chloroacetamide (20 mM, 30 min, RT) (Sigma-Aldrich, St Louis, USA). The reduced proteins were precipitated using Sera-Mag^TM^ carboxylate-modified magnetic beads (Cytiva, 24152105050250, Marlborough, USA) by adding ethanol (final concentration of 50%). The precipitated proteins were washed thrice with 80% ethanol following the manufacturer’s instructions. The precipitated proteins were then digested using trypsin (1:20 enzyme-to-substrate ratio) (Promega, Madison, USA) and incubated overnight at 37°C. The resulting peptides were precipitated using Sera-Mag^TM^ carboxylate-modified magnetic by adding acetonitrile (final concentration of ≥ 95%). The precipitated peptides were washed thrice with acetonitrile and eluted with 2% DMSO following the manufacturer’s instructions. Before the mass spectrometry acquisition, the samples were dried and resuspended in 0.1% formic acid (Buffer A) to achieve a final peptide concentration of 1 μg/3 μl. Aliquots from all replicates were pooled to ensure all peptides’ presence in the chromatogram library’s preparation.

### LC-Mass Spectrometry

Mass spectrometry was performed using an Orbitrap Fusion Lumos Tribrid Mass Spectrometer (Thermo Fisher Scientific, USA) coupled to a Dionex Ultimate 3000 UPLC system (Thermo Fisher Scientific). Peptides were loaded onto a PepMap 100 trap cartridge (0.3 x 5 mm, 5 μm C18, Thermo Fisher Scientific) and separated using an in-house 75 μm (inner diameter) analytical column with an integrated pulled tip emitter, packed with ReproSil-Pur 120 C18-AQ beads (1.9 μm, 120 Å, Dr Maisch, Ammerbuch, Germany) to 25 cm. For each injection, 1 μg of peptides were loaded and separated using a 120-min gradient from 3-31.2% buffer B (0.1% formic acid in 80% acetonitrile), followed by a 30-minute wash gradient and equilibration.

### Library generation and quantitative shotgun data-independent acquisition mass spectrometry (DIA-MS)

A pooled sample comprised of 1.5 μl of protein digest was used to generate a sample project-specific spectral library for data-dependent analysis (DDA). For each DDA analysis, 2.0 μl of the pooled sample was used with a 3-second cycle time instrument method. Liquid chromatography with a scan range of 350-1200 m/z was used. A narrow spectra (MS1) scan matching one of the six m/z mass ranges was performed using an Orbitrap resolution of 120,000. A normalised AGC target of 200% with an auto maximum injection time mode was used. An intensity threshold of 5.0e5 and dynamic exclusion duration of 60 seconds was employed for all data-dependent second stage (MS2) scans acquired at 30,000 orbitrap resolution, 400% normalised AGC target, 30% normalised HCD collision energy, with dynamic maximum injection time mode. For the DIA runs, the Orbitrap Fusion Lumos Tribrid Mass Spectrometer was configured to employ a series of variable-sized isolation windows for the MS2 fragmentation. An MS2 orbitrap resolution of 30,000, a normalised AGC target of 2000% with dynamic maximum injection time mode and normalised HCD collision energy of 30% were employed for all DIA scans. Precursor spectra over a 350-1200 m/z scan range were acquired before DIA scans with an orbitrap resolution of 120,000, normalised AGC target of 200%, and dynamic maximum injection time mode for all full scan MS spectra.

### ST3 Mfd Domain and Protein Structure Analysis

Given the impact of the *mfd* frameshift in yielding the hyperbiofilm mucoid phenotype of the isolates, we performed a more detailed in silico analysis of the Mfd protein integrating functional domain and structural predictions.

The *mfd* gene has been widely characterised in *E. coli* where it has been shown to have two functions: it dislodges RNA polymerase ternary elongation complexes through ATP hydrolysis and recruits the UvrABC excinuclease to repair sites of DNA damage, achieving this through binding to the UvrA subunit (Deaconescu et al., 2007; Selby and Sancar, 1993; 1995). The structure of Mfd in *E. coli* has been resolved experimentally (Supplementary Figure S7A) (Deaconescu and Darst, 2005).

The T1 *S. aureus mfd* gene is 1,168 AA long and is 100% identical to A0A9Q8DFZ8 Uniprot accession. It is 99.9% identical to the Uniprot accession Q2G0R8, only differing by a single amino acid (Asn1152Lys). Its domains have been characterised by InterPro including PFAM (Mistry et al., 2021) and SMART (Letunic *et al*., 2021) domain annotation homologous to the *E. coli mfd* gene. According to InterPro, the N-terminal region contains a domain homologous to the PFam PF17757 UvrB interaction domain relating to DNA-damage repair (139-226 AA) (Pakotiprapha et al., 2009; Paysan-Lafosse et al., 2023). It also contains a D3 domain homologous to PFAM PF21132 (402-468AA) (Kang et al., 2021), a CarD-like domain homologous to the SMART SM01058 (494-591 AA) (Nicolas et al., 1996), a DEAD/DEAH helicase domain homologous to PFAM PF00270 (de la Cruz et al., 1999) that is widely conserved across all domains of life (604-796 AA) and a C-terminal helicase domain homologous to PFAM PF00271 (808-969 AA). The domain at the C-terminal region is homologous to PF03461 (1020-1120 AA).

As no predicted structure was available for A0A9Q8DFZ8 in the Alphafold database (AFDB) (Varadi et al., 2022) (the AFDB accession of the extremely similar Q2G0R8 protein is AF-Q2G0R8-F1), we used Colabfold v1.5.3 webserver (Mirdita *et al*., 2022) to predict the structure of the T1 *mfd*.

The best model had a mean predicted local distance difference test (pLDDT) value of 83.85 and pTM (predicted template modelling score) of 0.67, indicating a well-modelled structural prediction overall (Supplementary Figure S7B). However, the predicted aligned error (PAE) matrix (Supplementary Figure S7D) suggests that the intra-module predictions are well-defined, but relative orientations between the modules in the structure are uncertain. Foldseek alignment against AFDB unsurprisingly confirmed that the entire *mfd* structure is well conserved across Gram-positive and Gram-negative bacteria, with good quality alignments across the entire structure (e-value < 1E-50 for homologs in *E. faecium*: AFDB accession AF-A0A132P774-F1-model_v4 and *Escherichia coli* K-12: AFDB accession AF-P30958-F1-model_v4).

The T3-T8 *mfd* gene product is also expected to yield a C-terminally truncated variant with 419 amino acids. Alignment with Foldseek revealed that it unsurprisingly had near perfect structural alignment across almost the whole protein (12-419AA) with the T1 *mfd* protein from residues 761-1168AA.

This indicates that the truncated protein retains only the C-terminal helicase and C-terminal domains losing modules relating to RNA polymerase binding and the DEAD/DEAH-box helicase domain.

Foldseek alignment against AFDB revealed that the C-terminal helicase domain (25-264AA in the truncated proteins) had strong alignment matches to the C-terminal domain of the ATP-dependent DNA helicase RecG (E-value 5.67E-14 against AFDB accession AF-P24230-F1-model_v4 for E coli).

The best model had a mean predicted pLDDT value of 76.07 and pTM (predicted template modelling score) of 0.45, indicating a decently modelled structural prediction overall (Supplementary Figure S7 C) but of lower quality than the full length Mfd protein. The PAE matrix (Supplementary Figure S7E) suggests that the two intra-domain predictions (i.e. the RecG like domain and the C-terminal domain) are well-defined, but relative orientations between them are uncertain.

## Notes

### Competing Interest Statement

The authors have declared no competing interest.

### Summary of Updates

All figure references in the main text of the manuscript have been fixed (fixing the Word -> PDF conversion)

